# Engineering inducible signaling receptors to enable erythropoietin-free erythropoiesis

**DOI:** 10.1101/2024.04.12.589260

**Authors:** Aadit P. Shah, Kiran R. Majeti, Freja K. Ekman, Sridhar Selvaraj, Eric Soupene, Prathamesh Chati, Roshani Sinha, Sofia E. Luna, Carsten T. Charlesworth, Travis McCreary, Benjamin J. Lesch, Tammy Tran, Devesh Sharma, Simon N. Chu, Matthew H. Porteus, M. Kyle Cromer

**Author notes:** These authors contributed equally to the study (Aadit P. Shah, Kiran R. Majeti).

## Abstract

Blood transfusion plays a vital role in modern medicine. However, availability is contingent on donated blood, and frequent shortages pose a significant healthcare challenge. *Ex vivo* manufacturing of red blood cells (RBCs) derived from universal donor O-negative pluripotent stem cells emerges as a solution, yet the high cost of recombinant cytokines required for *ex vivo* erythroid differentiation remains a major barrier. Erythropoietin (EPO) signaling through the EPO receptor is indispensable to RBC development, and EPO is one of the most expensive components in erythroid-promoting media. Here, we used design-build-test cycles to develop highly optimized small molecule-inducible EPO receptors (iEPORs) which were integrated at a variety of genomic loci using homology-directed repair genome editing. We found that integration of iEPOR at the endogenous *EPOR* locus in an induced pluripotent stem cell producer line enabled culture with small molecule to yield equivalent erythroid differentiation, transcriptomic changes, and hemoglobin production compared to cells cultured with EPO. Due to the dramatically lower cost of small molecules vs. recombinant cytokines, these efforts eliminate one of the most expensive elements of *ex vivo* culture media—EPO cytokine. Because dependence on cytokines is a common barrier to *ex vivo* cell production, these strategies could improve scalable manufacturing of a wide variety of clinically relevant cell types. More broadly, this work showcases how synthetic biology and genome editing may be combined to introduce precisely regulated and tunable behavior into cells, an advancement which will pave the way for increasingly sophisticated cell engineering strategies.

## Introduction

Blood cell transfusion plays an essential role in modern medicine. In support of surgery, obstetrics, trauma care, and cancer chemotherapy, approximately 35,000 units of blood are drawn daily in the U.S., contributing to an annual provision of 12 million red blood cell (RBC) units^1^. However, availability is contingent on donated blood, resulting in supply constraints and safety concerns. Blood shortages pose a significant global healthcare challenge, expected to worsen with aging populations and decreasing donor numbers^2^. Moreover, patient populations with especially rare blood types constitute up to 5% of blood transfusion cases^3^ and are most vulnerable to these shortages. From a financial perspective, the cost of RBC transfusion has been steadily increasing over the past two decades, accounting for nearly 10% of total inpatient hospital expenditure^4^. Collectively, these factors are expected to worsen the significant unmet medical need for transfusable blood.

To address these challenges, *ex vivo* manufacturing of RBCs in bioreactors from producer cell lines, such as pluripotent stem cells (PSCs), emerges as a renewable and scalable solution^5^. Early clinical trials have shown that *ex vivo*-derived RBCs may be delivered to patients with no reported adverse events^6^. In addition, *ex vivo* -derived RBCs offer potential benefits compared to donor blood, including a lower risk of infectious disease transmission, streamlined production, product uniformity, and ability to source or genetically engineer antigen-negative cells^2^. However, *ex vivo* RBC production is still prohibitively expensive, owing in large part to the high cost of recombinant cytokines required to stimulate producer cells to expand and differentiate into erythroid cells^7^. Erythropoietin (EPO) signaling through the EPO receptor (EPOR) is indispensable to RBC development^8^, and of all components in erythroid-promoting media, EPO is one of the most expensive^7^. Given prior success manipulating the EPOR to increase erythropoietic output^9^ and the ease with which erythroid development is modeled *ex vivo*^10^, in this work we used synthetic biology tools and genome editing technology to de-couple EPOR signaling from the EPO cytokine.

The cellular mechanisms that regulate erythroid differentiation from hematopoietic stem and progenitor cells (HSPCs) are well understood, and efficient differentiation requires activation of the EPOR/JAK/STAT signaling cascade by EPO^11^. In its native form, two EPOR monomers dimerize in the presence of EPO to activate downstream signaling^12^. Prior work has shown that EPOR dimerization may be initiated by a range of dimer orientations and proximities using agonistic diabodies or in the context of chimeric receptors^12–14^. Because mutant FK506 binding proteins (FKBP)-based dimerization domains have been deployed to create small molecule-inducible safety switches^15^, we hypothesized that FKBP domains could be repurposed to create synthetic EPOR receptors to place EPO signaling under control of a small molecule. Here, we demonstrate that EPOR signaling can be induced by small molecule stimulation of highly optimized chimeric receptors—hereafter termed inducible EPORs (iEPORs). We then used homology-directed repair genome editing to integrate these iEPORs under regulation of various endogenous and exogenous promoters to identify strategies that best recapitulate native EPOR signaling.

This work establishes iEPORs as a tool that enables highly efficient *ex vivo* production of RBCs using a low-cost small molecule. By removing dependence on one of the most expensive elements of *ex vivo* erythrocyte production, these efforts address one of the major barriers to meeting the global demand for blood with *ex vivo* -manufactured RBCs. More broadly, this work demonstrates how synthetic biology and genome editing may be combined to introduce precisely regulated and tunable behavior into cells for a wide variety of therapeutic applications.

## Results

### FKBP-EPOR chimeras enable small molecule-dependent erythropoiesis

To determine whether FKBP domains could be successfully repurposed to dimerize EPOR monomers and initiate downstream EPOR signaling, we first designed a set of seven candidate FKBP-EPOR chimeras. These included placement of the FKBP domain at the N-terminus, C-terminus, at various locations within the native EPOR, and as a full replacement of the EPOR extracellular domain (**Fig. 1A** ). DNA donor templates corresponding to each design were packaged in AAV6 vectors and integrated into the *CCR5* safe harbor site in human primary HSPCs using combined CRISPR/AAV6-mediated genome as previously described^16–18^. Expression of each FKBP-EPOR chimera was driven by the strong, constitutive SFFV promoter followed by a 2A-YFP to allow fluorescent readout of edited cells (**Fig. 1A** ). Edited HSPCs were then subjected to an established 14-day *ex vivo* erythrocyte differentiation protocol^19,20^ in the absence of EPO and with or without 1nM of FKBP dimerizer AP20187 small molecule (hereafter referred to as “BB” dimerizer)^15^. Since EPO is essential for differentiation, we hypothesized that erythroid differentiation would only occur when BB stimulated a functional iEPOR to activate downstream signaling (**Fig. 1B** ).

**Figure 1.**
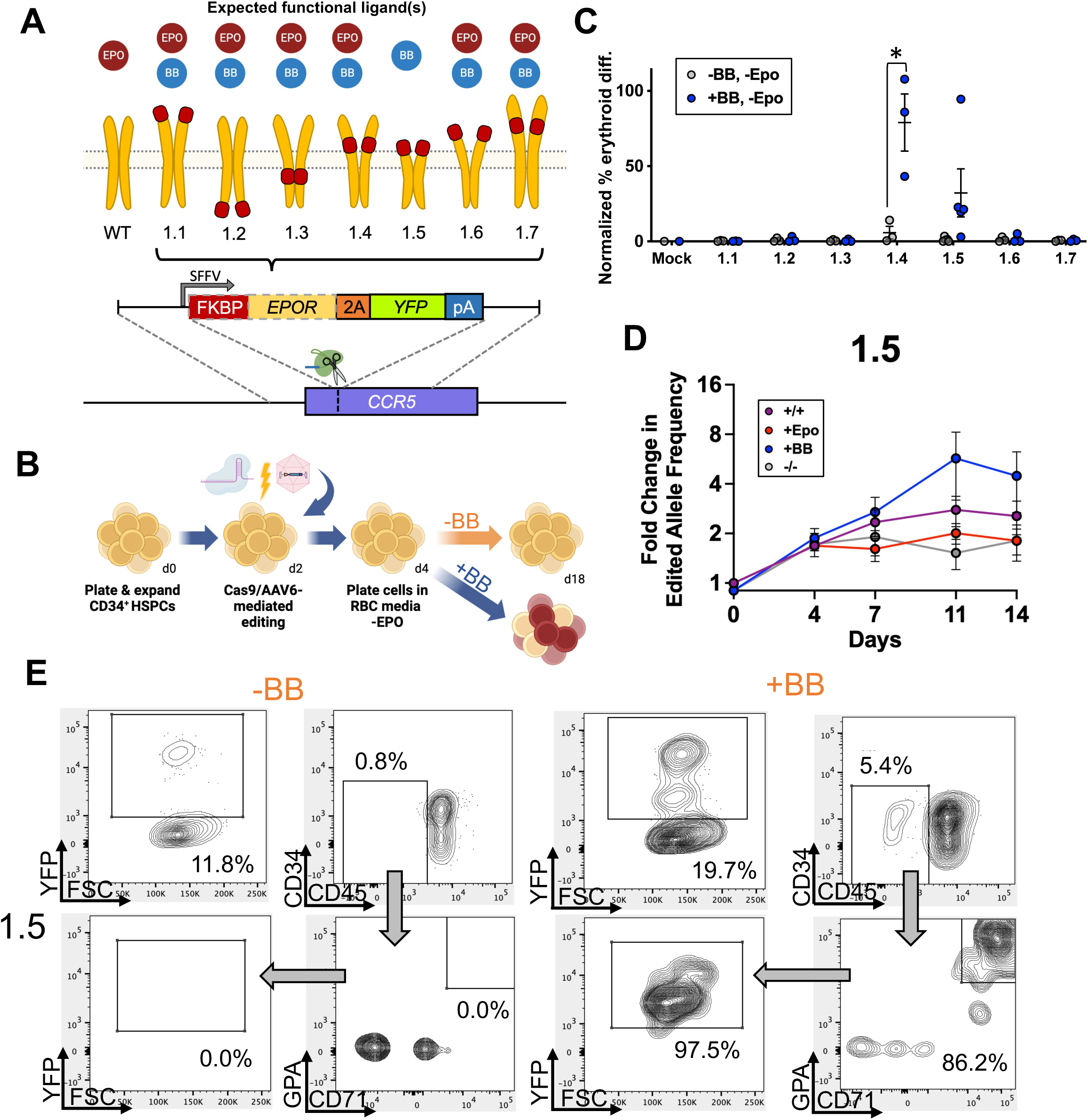
**Screening of FKBP-EPOR chimeras to facilitate EPO-free erythroid differentiation.** A: Schematic of chimeric FKBP-EPOR transgenes integrated at the *CCR5* locus via CRISPR/AAV-mediated editing. Red boxes represent the location of FKBP within the EPOR. B: Schematic of HSPC editing and subsequent erythroid differentiation. C: Percentage of edited HSPCs that acquired erythroid markers (CD34^-^/CD45^-^/CD71^+^/GPA^+^) +/-BB normalized to unedited cells +EPO at d14 of differentiation. Bars represent median +/-SEM; * = p<0.05 by unpaired t-test. D: Percent edited alleles over the course of differentiation +/-BB and +/-EPO. Bars represent median +/-SEM. E: Representative flow cytometry staining and gating scheme for iEPOR 1.5-edited HSPCs at d14 of differentiation -EPO and +/-BB. Arrows indicate that only gated cells are displayed on the subsequent plot.

At the end of differentiation, we stained cells for established erythroid markers and analyzed by flow cytometry (**Supplementary Fig. S1** ). As expected, we found that unedited “Mock” conditions yielded no erythroid cells (CD34^-^/CD45^-^/CD71^+^/GPA^+^), while HSPCs edited with iEPOR designs 1.4 and 1.5 showed BB-dependent erythroid differentiation (**Fig. 1C** ; **Supplementary Fig. S2** ). Although FKBP-EPOR design 1.4 appeared to be most effective, for downstream optimizations we iterated on design 1.5 due to the smaller cassette size and because removal of the entire EPOR extracellular domain is expected to eliminate potential activation by EPO cytokine. This allowed us to create a receptor that could activate the EPOR pathway only when dimerizer was present but not when endogenous hormone was present. Further investigation of iEPOR 1.5 found a >4x selective advantage imparted to edited cells by the end of erythroid differentiation when cells were cultured in the presence of BB without EPO as indicated by increasing edited allele frequency measured by droplet digital PCR (ddPCR)(**Fig. 1D**). In addition, virtually all cells that acquired erythroid markers in the iEPOR 1.5 condition were YFP^+^ (**Fig. 1E** ), indicating that only edited cells were capable of differentiation.

To investigate why certain FKBP-EPOR designs were non-functional, we used AlphaFold2^21^ to generate *in silico* structure predictions of each candidate iEPOR in comparison to wild-type EPOR. We observed a high-confidence structure generated across wild-type EPOR extracellular and transmembrane domains, with low-confidence scores given to signal peptide and intracellular regions (**Supplementary Fig. S3** ). For candidate iEPORs, we observed a high-confidence structure corresponding to the FKBP domain at the anticipated location among all designs. Although this analysis did not reveal any obvious protein structure disruption caused by addition of FKBP domains to the EPOR protein, our experiments demonstrated that FKBP placement within the EPOR has a great bearing on its signaling potential. We found that only those constructs with FKBP placed immediately upstream of the EPOR transmembrane domain could initiate BB-dependent signaling. Therefore, it is possible that designs with FKBP within the intracellular domain may interfere with JAK/STAT signaling, while FKBPs placed further upstream of the transmembrane domain may not mediate sufficient proximity of EPOR intracellular domains to achieve sustained signaling.

### Signal peptides and hypermorphic EPOR mutation increase iEPOR potency

Initial iEPOR designs 1.4 and 1.5 both mediated BB-dependent erythroid production, yet they were unable to achieve a level of differentiation equivalent to unedited cells cultured with EPO (mean of 78.9% and 32.2% of the amount of differentiation achieved with unedited cells +EPO for iEPOR 1.4 and 1.5, respectively; **Fig. 1C** ). Therefore, we engineered second-generation iEPORs to determine if addition of a signal peptide (SP) onto iEPOR 1.5 could enhance potency, since elimination of the entire EPOR extracellular domain also removes the native SP at the N-terminus. To test the effect of these modifications, we designed and built constructs that added the native EPOR SP or the IL6 SP^22^ onto the N-terminus of iEPOR 1.5. This comparison was performed because SPs for cytokines are known to be particularly strong^23,24^, and we observed in unpublished work that the IL6 SP effectively mediates export to the HSPC membrane (data not shown). These DNA donor templates were packaged into AAV6 and integrated into the *CCR5* locus as before (**Fig. 2A** ). We then performed *ex vivo* erythroid differentiation in the presence or absence of EPO and BB. We found that addition of EPOR SP and IL6 SP both improved mean erythroid differentiation in the presence of BB alone (44.5% and 62.6%, respectively) compared to the original iEPOR 1.5 design (32.2%)(**Fig. 2B** & **Supplementary Fig. S4** ). These vectors also yielded a further selective advantage in the presence of BB (both with and without EPO), achieving a mean 10.0- and 10.6-fold increase in edited allele frequency by the end of erythroid differentiation with addition of EPOR and IL6 SPs, respectively (**Fig. 2C** & **Supplementary Fig. S5A** ).

**Figure 2:**
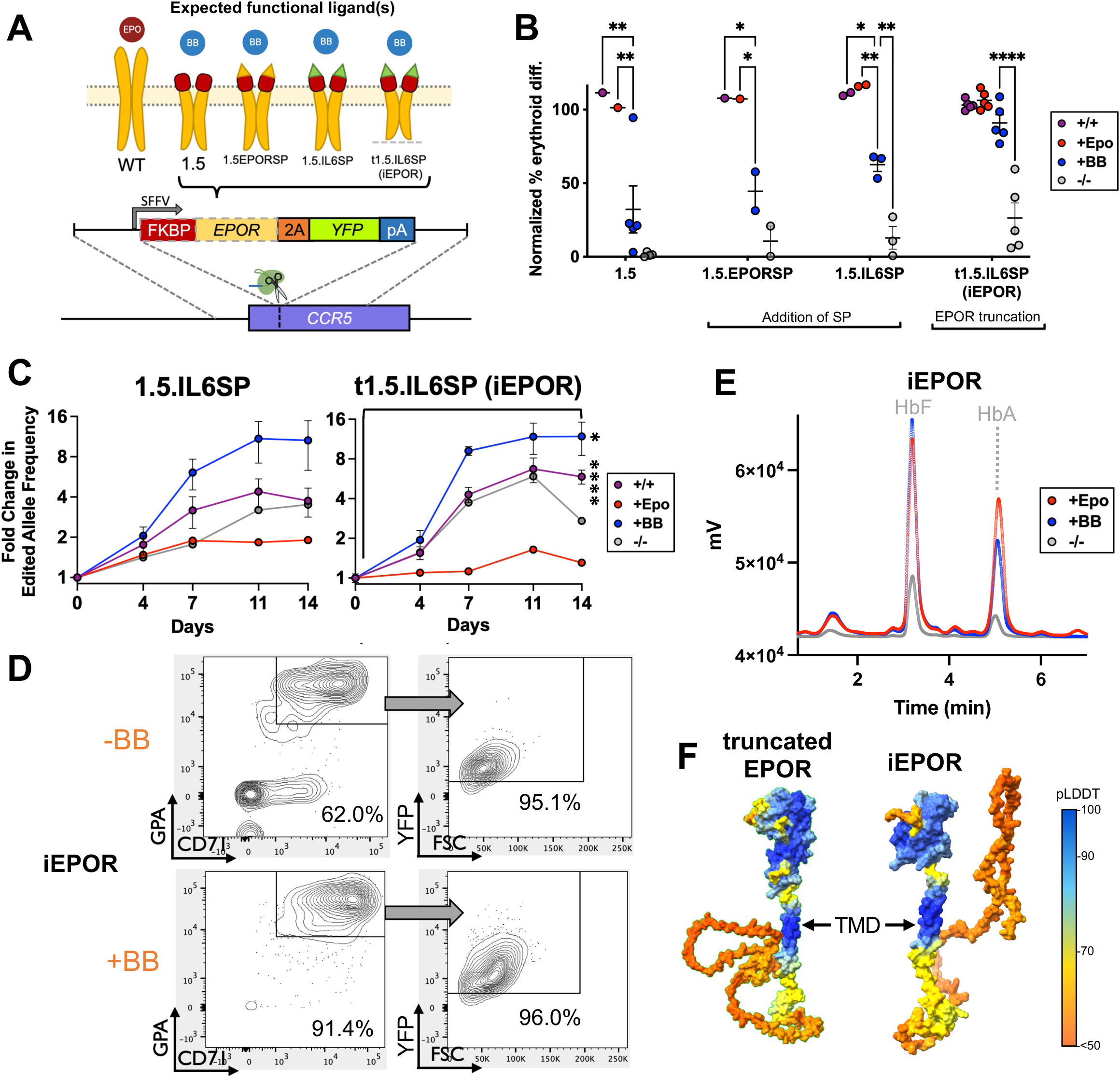
**Modulation of iEPOR effect by addition of signal peptide & EPOR truncation.** A: Schematic of second-generation iEPORs integrated at *CCR5* locus. Red boxes represent the FKBP domain; yellow and green triangles indicate EPOR and IL6 SPs, respectively; dashed line represents EPOR truncation. B: Percentage of edited HSPCs that acquired erythroid markers (CD34^-^/CD45^-^/CD71^+^/GPA^+^) +/-BB normalized to unedited cells +EPO at d14 of differentiation. iEPOR 1.5 data from Fig. 1C shown for comparison. Bars represent median +/-SEM; * = p<0.05, ** = p<0.01, and **** = p<0.0001 by unpaired t-test. C: Percent edited alleles over the course of differentiation +/-BB and +/-EPO. Bars represent median +/-SEM; * = p<0.05 and **** = p<0.0001 comparing d0 vs. d14 within treatment by unpaired t-test. D: Representative flow cytometry staining and gating scheme for iEPOR-edited HSPCs at d14 of differentiation -EPO and +/- BB. Arrows indicate that only gated cells are displayed on the subsequent plot. E: Representative hemoglobin tetramer HPLC plots at d14 of erythroid differentiation. +BB and -BB/-EPO conditions were from cells edited with iEPOR; +EPO condition was from unedited cells. All plots normalized to 1e6 cells. F: AlphaFold2-based structure prediction of truncated EPOR and iEPOR. SP was removed since this sequence will be cleaved following translocation to the membrane. TMD labeled with an arrow as a reference point.

Given the higher efficacy of iEPOR 1.5 with IL6 SP, we investigated whether incorporation of a naturally occurring nonsense mutation (*EPOR* ^W439X^) that truncates the 70 C-terminal amino acids of EPOR and eliminates a negative inhibitory domain may additionally increase receptor potency^9^. Therefore, we designed a vector with this truncated EPOR intracellular domain as well as IL6 SP and observed a further enhancement, achieving a mean of 90.9% erythroid differentiation compared to EPO-cultured HSPCs (**Fig. 2B** ). This significantly increased the selective advantage of edited cells cultured in the presence of BB, achieving a mean 11.9-fold increase in edited alleles by the end of erythroid differentiation (**Fig. 2C** ). As before, virtually all cells that acquired erythroid markers were YFP^+^, indicating that only edited cells stimulated with BB were able to initiate EPOR signaling (**Fig. 2D** ). Notably, a substantial portion of cells also differentiated in the absence of BB, which we addressed in downstream experiments. We will hereafter refer to our optimized FKBP-EPOR design 1.5 with IL6 SP and naturally occurring truncation as “iEPOR”).

To ensure that iEPOR-stimulated erythroid cells produce functional hemoglobin, we performed hemoglobin tetramer high-performance liquid chromatography (HPLC) at the end of erythroid differentiation. We found that cells edited with the optimized iEPOR and cultured with BB yielded a hemoglobin production profile consisting primarily of adult and fetal hemoglobin (HbA and HbF, respectively). This hemoglobin production profile was indistinguishable from that produced by unedited cells culture with EPO (**Fig. 2E** ).

Finally, we used AlphaFold2 to predict the structure of the optimized iEPOR and find remarkable similarity to the predicted structure of the naturally occurring truncated EPOR (**Fig. 2F**). As expected, we observe a shortening of the low-confidence intracellular domain for the truncated EPOR compared to wild-type EPOR (**Supplementary Fig. S6** ) as well as a high-confidence structure corresponding to the FKBP domain in the expected location for the optimized iEPOR. As with native EPOR SP, we also observe a low-confidence region corresponding to the IL6 SP.

### iEPOR expression profile impacts receptor function

While our optimized iEPOR was effective at mediating small molecule-dependent erythroid differentiation and hemoglobin production, we observed some erythroid differentiation and hemoglobin production in the absence of BB as well (**Fig. 2B-E** ). This could be due to the strong, constitutive viral SFFV promoter driving supraphysiologic levels of receptor expression that induced ligand-independent dimerization. In contrast to the potent SFFV promoter, prior work has shown that CD34^+^ HSPCs express low levels of the endogenous *EPOR* , and expression increases modestly over the course of *ex vivo* erythroid differentiation^25^. Therefore, in the next round of optimizations, we explored the impact of various expression profiles on iEPOR activity. To do so, we developed targeted integration strategies that placed an identical optimized iEPOR under expression from: 1) an exogenous yet weaker, constitutive human *PGK1* promoter following integration at the *CCR5* locus (hereafter referred to as “*PGK* (iEPOR)”); 2) the strong erythroid-specific *HBA1* promoter following integration into the start codon of the *HBA1* locus^10^ (hereafter referred to as “*HBA1*(iEPOR)”); and 3) the endogenous *EPOR* locus following integration into the 3’ end of the gene and linked by a 2A cleavage peptide (hereafter referred to as “*EPOR* (iEPOR)”)(**Fig. 3A** ). We chose these additional integration strategies to investigate whether extremely high *iEPOR* expression or simply constitutive expression throughout differentiation was most responsible for the dimerizer-independent activity of SFFV(iEPOR). These experiments also investigated whether erythroid-specific expression of iEPOR from the highly expressed *HBA1* locus may elicit the most dramatic pro-erythroid effect or if, alternatively, integration of iEPOR at the endogenous *EPOR* locus may best recapitulate native EPOR signaling—analogous to the effective regulation of synthetic T cell receptors when knocked into the native *TRAC* locus^26^.

**Figure 3:**
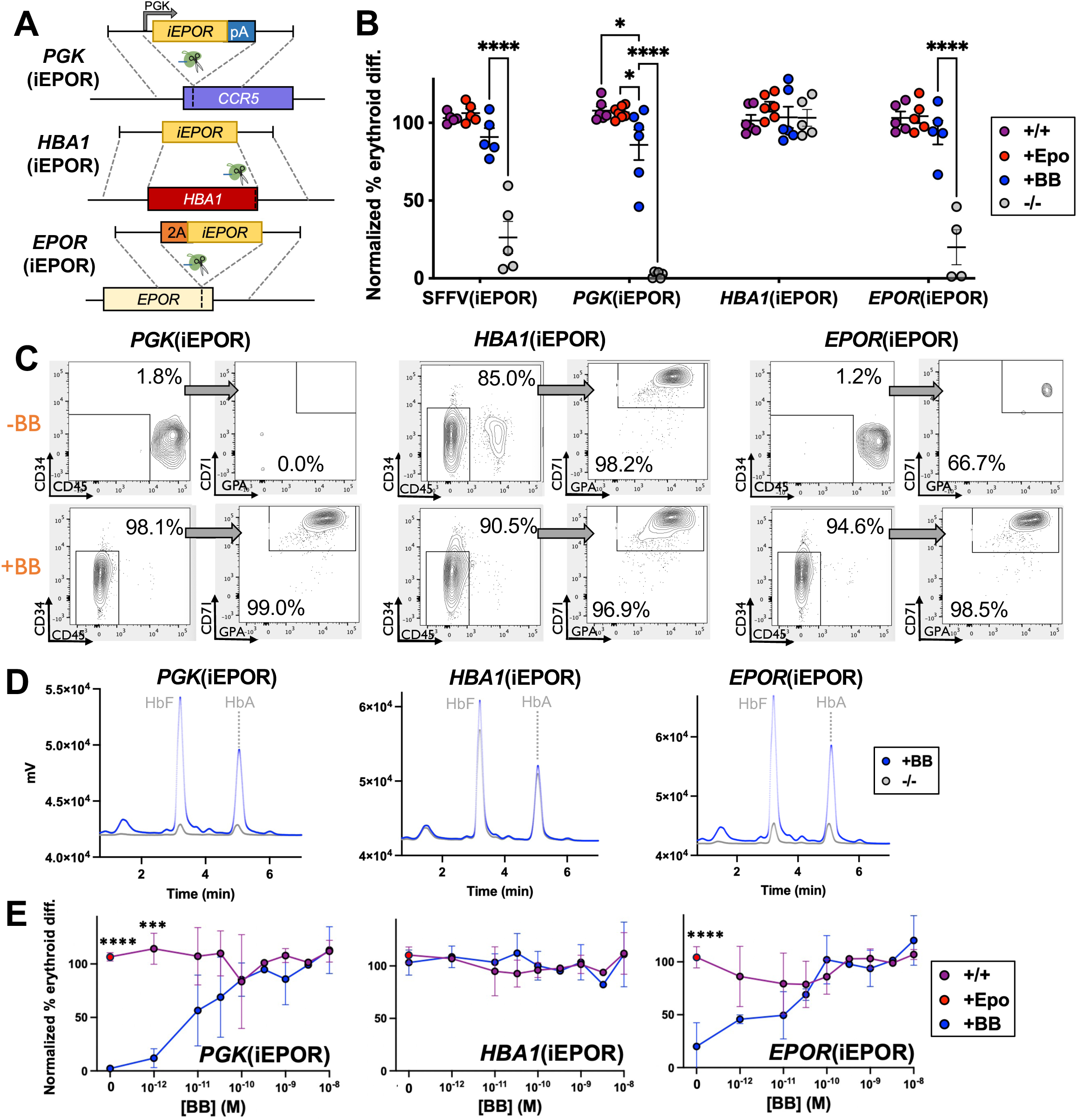
**Modulation of iEPOR effect by expression from various promoters.** A Schematic of third-generation iEPORs that drive expression from: 1) PGK promoter from *CCR5* safe harbor site; 2) erythroid-specific *HBA1* locus; and 3) endogenous *EPOR* locus. B: Percentage of edited HSPCs that acquired erythroid markers (CD34^-^/CD45^-^/CD71^+^/GPA^+^) +/-BB normalized to unedited cells +EPO at d14 of differentiation. SFFV(iEPOR) data from Fig. 2B shown here for comparison. Bars represent median +/-SEM; * = p<0.05, and **** = p<0.0001 by unpaired t-test. C: Representative flow cytometry staining and gating scheme for edited HSPCs at d14 of differentiation -EPO and +/-BB. Arrows indicate that only gated cells are displayed on the subsequent plot. D: Representative hemoglobin tetramer HPLC plot of edited HSPCs at d14 of differentiation -EPO and +/-BB. E: Dose response of edited HSPCs cultured over a range of [BB] at d14 of differentiation normalized to unedited cells +EPO. Bars represent median +/-SEM; *** = p<0.001 and **** = p<0.0001 comparing +EPO/+BB to +BB conditions by unpaired t-test.

Following integration of each vector into the intended site in primary HSPCs, we performed *ex vivo* erythroid differentiation in presence or absence of EPO and BB. We observed that all three integration strategies yielded effective erythroid differentiation in presence of BB compared to unedited cells cultured with EPO (**Fig. 3B** & **C**). However, the greatest differences were found in edited conditions cultured without BB or EPO. Compared to the mean 26.3% erythroid differentiation we observed previously in the SFFV(iEPOR)-edited condition without EPO or BB, expression of iEPOR from the *PGK* and *EPOR* promoters both reduced BB-independent activity (mean of 2.2% and 20.0% in *PGK* (iEPOR) and *EPOR* (iEPOR) conditions, respectively)(**Fig. 3B** & **C**). In contrast, we found that expression of iEPOR from the *HBA1* promoter drove high frequencies of erythroid differentiation in the presence and absence of BB, indicating constitutive activity (**Fig. 3B** & **C**). Because *HBA1* is expressed much more highly than *EPOR* by the end of *ex vivo* differentiation^27^, we hypothesize this BB-independent activity could be a result of supraphysiologic levels of *iEPOR* expression from the *HBA1* promoter that leads to spontaneous signaling even in absence of dimerizing ligand. As before, we confirmed that each iEPOR integration strategy yielded normal production of adult and fetal hemoglobin when edited cells were cultured in the presence of BB without EPO (**Fig. 3D** ). Due to the high level of erythroid differentiation observed in the *HBA1*(iEPOR) condition, it was unsurprising that edited cells cultured with neither EPO nor BB also produced a substantial amount of adult and fetal hemoglobin.

Next, we determined whether expression of *iEPOR* from these different promoters has a bearing on the dose response to BB. While prior work using BB found 1nM to be most effective at activating small molecule-inducible safety switches^15^, we observed substantial erythroid differentiation at levels well below 1nM of BB. In fact, we found that 1pM and 10pM of BB yielded erythroid differentiation that was comparable to EPO in cells edited with *EPOR* (iEPOR) and *PGK* (iEPOR) strategies, respectively (**Fig. 3E** ). However, to achieve mean differentiation that was identical to or greater than EPO-stimulated cells required a dose of 0.1nM for *PGK* (iEPOR)- and *EPOR* (iEPOR)-edited populations. In contrast, we found that cells edited with *HBA1*(iEPOR) yielded efficient erythroid differentiation across the entire dose range, including in the absence of BB (**Fig. 3E** ), consistent with constitutive activity of this integration strategy.

### iEPOR closely replicates native EPOR signaling

In its native form, EPO cytokine dimerizes two EPOR monomers, leading to a JAK/STAT signaling cascade culminating in translocation of phosphorylated STAT5 to the nucleus, which initiates a pro-erythroid transcriptional program^11^. While we have shown that iEPOR-edited cells stimulated with BB acquire classic erythroid markers and produce normal hemoglobin profiles, an open question is whether this synthetic stimulus recapitulates the complex transcriptional response of native EPOR signaling (**Fig. 4A** ). To investigate this, we edited HSPCs with our various iEPOR integration strategies (**Fig. 3A** ) and performed bulk RNA-sequencing (RNA-Seq) on at d14 of erythroid differentiation in absence of EPO and presence of BB. For comparison, we also performed RNA-Seq on unedited cells at the beginning (d0) and end (d14) of erythroid differentiation in the presence of EPO. These efforts yielded an average of 55.1M reads per sample with 98.5% of reads aligned to the genome and 97.2% with Quality Score ≥20 (**Supplementary Fig. S7** ).

**Figure 4:**
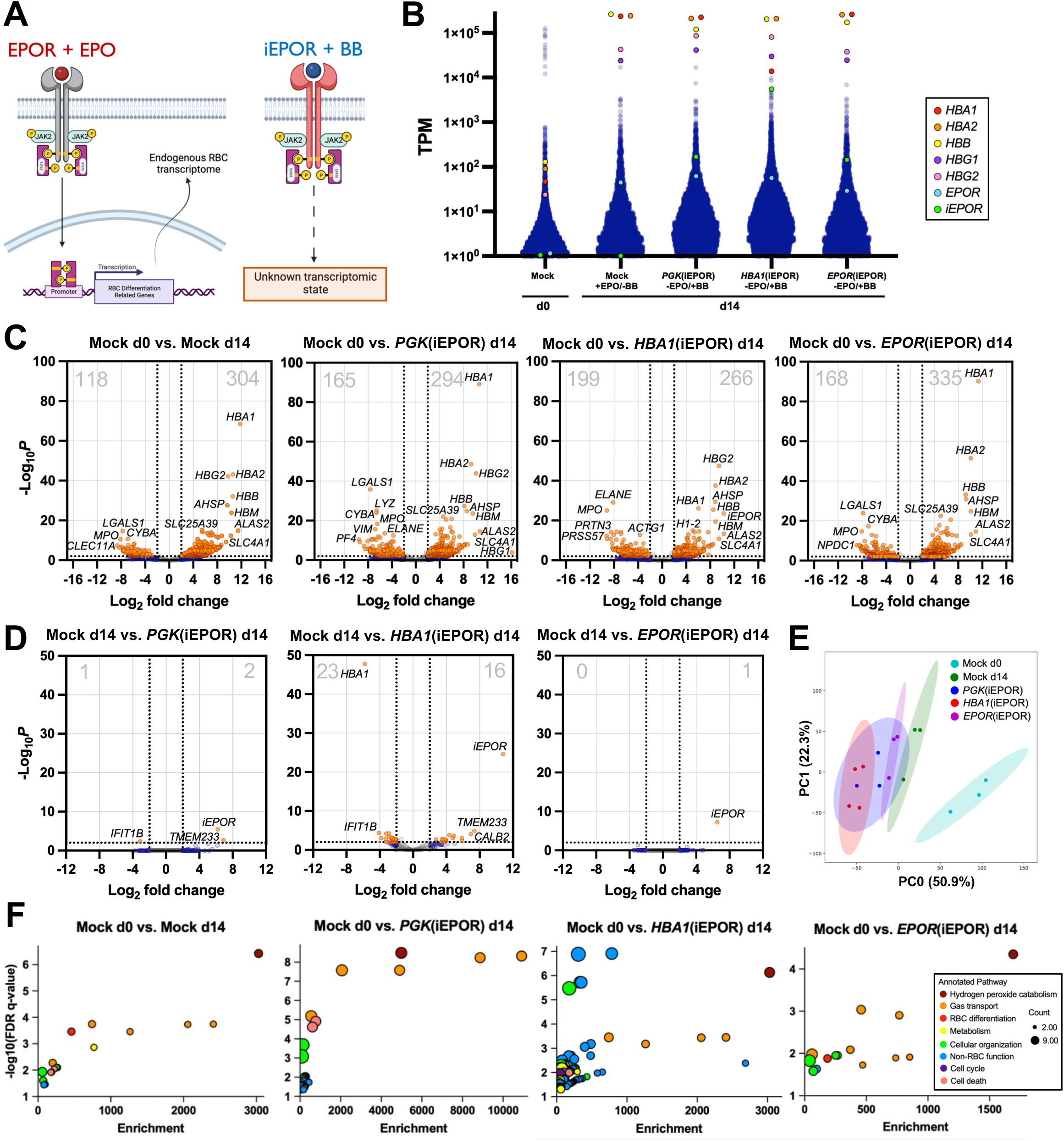
**Transcriptome-wide comparison of iEPOR-edited cells to EPO-differentiated cells.** A: Schematic of well-characterized native EPO+EPOR signaling effects vs. undefined BB+iEPOR signaling effects. B: Transcripts per million (TPM) from RNA-Seq with annotation for globin, *EPOR*, and *iEPOR* genes. C: Volcano plot comparing unedited and edited HSPCs at d14 of differentiation vs. unedited HSPCs at d0. Dashed lines are drawn at +/-2 log_2_ fold change and p=0.01. Total number of significantly down- and upregulated genes is shown in top left and top right of each plot, respectively. D: Volcano plot comparing edited HSPCs at d14 +BB vs. unedited HSPCs at d14 +EPO. Dashed lines are drawn at +/-2 log_2_ fold change and p=0.01. Total number of significantly down- and upregulated genes is shown in top left and top right of each plot, respectively. E: Principal component analysis of all conditions with covariance ellipses. F: Summary of gene ontology (GO) analysis comparing all d14 conditions vs. d0 control. Differentially expressed genes from volcano plots (Fig. 4C) were used as input. Specific GO pathways were binned into broader categories and enrichment score was derived by Enrichr software. Count refers to the number of genes within each GO pathway that contributed to enrichment.

In analyzing these data, we found that alpha-, gamma-, and beta-globin are among the most significantly upregulated genes in unedited cells at d14 vs. d0 (**Fig. 4B** & **C**). Similarly for all cells edited with iEPOR, these globins are also among the most significantly upregulated genes (**Fig. 4B** & **C**). In fact, by the end of differentiation these globins comprise a mean of 86.5%, 76.9%, 63.3%, and 83.7% of all reads for unedited cells cultured with EPO as well as *PGK* (iEPOR)-, *HBA1*(iEPOR), and *EPOR* (iEPOR)-edited cells cultured with BB, respectively (**Supplementary Fig. S8A** ). As previously observed, we found that *EPOR* expression increases over the course of erythroid differentiation^25^ (38.8-fold from d0 to d14 in unedited cells; **Fig. 4B** & **Supplementary Fig. S8B** ). In all iEPOR-edited cells we observe roughly equivalent levels of *EPOR* compared to unedited cells, which is expected since each integration strategy preserves native *EPOR* expression. As for *iEPOR* expression, we find that both *PGK* and *EPOR* promoters drive expression comparable to that of native *EPOR* at d14 in unedited cells (**Fig. 4B** & **Supplementary Fig. S8C** ). However, the *HBA1* promoter drives supraphysiologic levels of *iEPOR* , with expression nearing that of the globins. Since the *HBA1*(iEPOR) integration strategy replaces a full copy of the *HBA1* gene with *iEPOR* transgene, it is not surprising to find a significant decrease in *HBA1* expression in this condition as well (**Fig. 4B** , **Supplementary Fig. S8A** , & **S8D** ). Consistent across donors, genes most highly expressed in HSPCs are uniformly downregulated in all d14 samples while erythroid-specific genes are uniformly upregulated (**Fig. 4B** , **4C**, & **Supplementary Fig. S8D-G** ). Because of this, we find that d0 HSPCs and all d14 samples segregate into two distinct hierarchies (**Supplementary Fig. 9A** ), indicating a high degree of similarity across all d14 samples regardless of whether these were unedited cells cultured with EPO or iEPOR-edited cells cultured with BB.

Although consistent differences were observed comparing all conditions to d0 HSPCs, we next determined whether significant differences existed at the end of differentiation between unedited cells cultured with EPO and iEPOR-edited conditions cultured with BB. This comparison revealed an extremely high degree of similarity between unedited cells cultured with EPO and *PGK* (iEPOR)-edited cells cultured with BB; only three genes were differentially expressed, including upregulation of the *iEPOR* transgene (**Fig. 4D** ). In contrast, the transcriptome of *HBA1*(iEPOR)-edited cells departed more substantially from unedited cells, with a total of 47 differentially expressed genes. As expected, in this condition we observed significant upregulation of *iEPOR* as well as downregulation of *HBA1*. Remarkably, we find that the only differentially expressed gene in *EPOR* (iEPOR)-edited conditions is the *iEPOR* transgene, indicating that this condition best recapitulated native EPOR signaling. These conclusions were further supported by principal component analysis, which found that all d14 samples clustered separately from d0 samples and that *EPOR* (iEPOR)-edited cells stimulated with BB most closely resemble unedited cells cultured with EPO (**Fig. 4E** ). Gene co-expression network analysis additionally revealed a high degree of similarity between iEPOR-edited conditions and unedited cells cultured with EPO (**Supplementary Fig. S9B** ).

To determine which cellular processes were activated by EPO compared to edited cells cultured with BB, we performed gene ontology analysis of differentially expressed genes in each condition compared to unedited cells at d0 (**Fig. 4F** ). At d14, the most highly enriched pathways were hydrogen peroxide (H_2_O_2_) catabolism—a critical function of erythrocytes to process the significant amounts of superoxide and H O that occur during oxygen transport^28^. We also find gas transport and erythroid differentiation processes to be highly enriched across all d14 samples. From this analysis, the *HBA1*(iEPOR) condition shows the most substantial departure from native EPOR signaling, with a number of significantly enriched pathways unrelated to erythrocyte function. On the other hand, we find that *EPOR* (iEPOR) most closely resembles native EPOR signaling, leading us to conclude that expression of synthetic receptors from the endogenous promoter is likely to best recapitulate the transcriptomic changes initiated by native cytokine signaling.

### iEPOR enables EPO-free erythropoiesis from O-negative induced pluripotent stem cells

All prior work was done in primary HSPCs to determine whether we could successfully engineer small molecule-inducible EPORs that recapitulate native erythroid development and function. However, while primary hematopoietic HSPCs may be sourced from umbilical cord blood and mobilized peripheral blood to produce RBCs *ex vivo* , their expansion capacity is limited^2^. As a solution, induced pluripotent stem cell (iPSC) producer lines provide a potentially unlimited source of patient-derived RBCs^6^. Therefore, in downstream experiments we used an iPSC line called PB005 derived from a healthy donor with O-negative blood type^29^ to determine if iEPORs could effectively produce erythroid cells from a “universal” blood donor.

To test this, we integrated our most effective *iEPOR* expression strategies— *PGK* (iEPOR) and *EPOR* (iEPOR)—into the PB005 iPSC line and isolated homozygous knock-in clones (**Fig. 5A** & **Supplementary Fig. S10A** ). These clones were then subjected to an established 12-day differentiation into hematopoietic progenitor cells (HPCs). Surprisingly, we found that *EPOR* (iEPOR)-edited clones yielded a substantially greater total cell count compared to both unedited and *PGK* (iEPOR)-edited clones (**Supplementary Fig. S10B** ), although this condition had a slightly higher proportion of cells staining for erythroid markers (**Supplementary Fig. S10C & D** ). Following iPSC-to-HPC differentiation, we performed a 14-day RBC differentiation without EPO and +/-BB (**Fig. 5A** ). We found that all cells (unedited and edited) effectively differentiated in the presence of EPO, whereas no erythroid differentiation was observed in unedited cells in the absence of EPO (**Fig. 5B** & **Supplementary Fig. S11A** ). *PGK* (iEPOR)-edited cells stimulated with BB yielded a high percentage of erythroid cells, but differentiation efficiency was significantly less than that achieved by EPO in these clones at every timepoint (**Fig. 5B** ). In addition, overall cell proliferation was substantially lower than that achieved with EPO (**Fig. 5C** ). In contrast, *EPOR* (iEPOR) clones achieved a differentiation efficiency that was indistinguishable from clones cultured with EPO; cell proliferation over the course of differentiation was also nearly equivalent to that achieved with EPO (**Fig. 5B** & **C**). Given frequent clonal differences observed in proliferation capacity, we also examined cell proliferation from the best *PGK* (iEPOR)- and *EPOR* (iEPOR)-edited clones. By day 14, the most highly proliferative *PGK* (iEPOR)-edited clone only achieved 37.3% of the proliferation of that same clone when cultured with EPO (**Supplementary Fig. S11B** ). However, the most effective *EPOR* (iEPOR)-edited clone achieved even greater proliferation (107.8%) compared to the same clone cultured with EPO.

**Figure 5:**
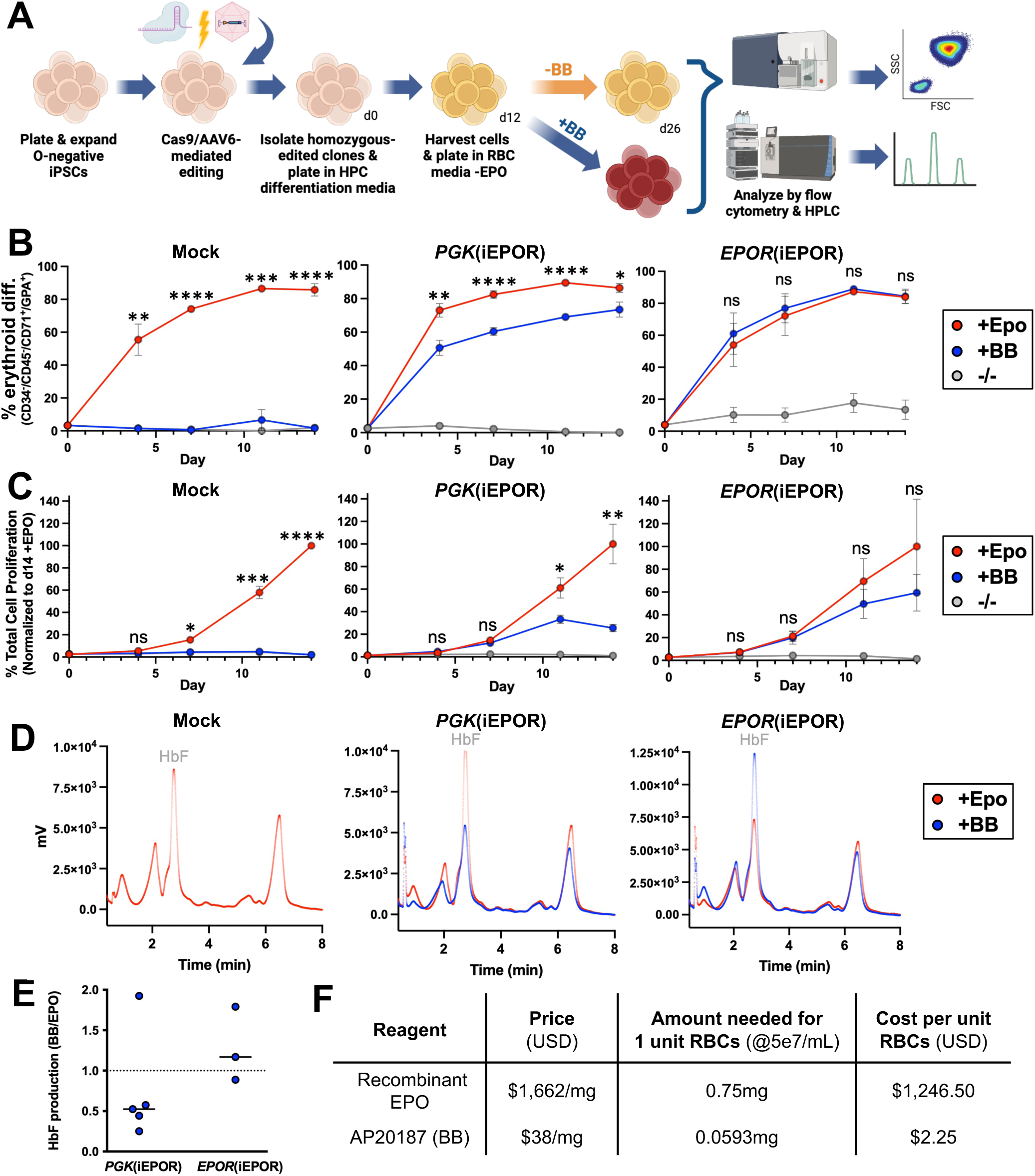
**Differentiation of iPSCs into erythroid cells using iEPOR+BB compared to exogenous EPO.** A: Schematic of iPSC-to-erythroid cell differentiation strategy and subsequent analysis. B: Percentage of cells that acquired erythroid markers (CD34^-^/CD45^-^/CD71^+^/GPA^+^) over the course of differentiation. Bars represent mean +/-SEM; ns = not statistically significant, * = p<0.05, ** = p<0.01, *** = p<0.001, and **** = p<0.0001 comparing +BB to +EPO conditions by unpaired t-test. C: Percentage of total cell proliferation normalized to clones cultured +EPO over the course of differentiation. Bars represent mean +/SEM; ns = not statistically significant, * = p<0.05, ** = p<0.01, *** = p<0.001, and **** = p<0.0001 comparing +BB to +EPO conditions by unpaired t-test. D: Representative hemoglobin tetramer HPLC plots of edited and unedited iPSC-derived erythroid cells at end of differentiation. E: Ratio of HbF production in +BB vs. +EPO conditions of iEPOR-edited iPSC-derived erythroid cells at end of differentiation. F: Cost comparison of EPO and BB (lowest price per mg commercially available for purchase as of 2/9/24).

Next, we measured the hemoglobin profiles of these iPSC-derived erythroid cells using HPLC and observed fetal hemoglobin to be the most prevalent tetramer in the presence of EPO, which is consistent with prior studies^30^. We found this to be the case as well for clones edited with both *PGK* (iEPOR) and *EPOR* (iEPOR) conditions cultured with BB (**Fig. 5D** ). While *PGK* (iEPOR) conditions cultured with BB almost uniformly expressed lower fetal hemoglobin than their EPO-cultured counterparts, we observed the opposite for *EPOR* (iEPOR)-edited conditions, with clones cultured with BB typically producing elevated levels of fetal hemoglobin relative to those same clones cultured with EPO (**Fig. 5D** & **E**). However, there appeared to be some clonal variation since not all clones conformed to these trends (**Fig. 5E** ). These findings were further confirmed by quantifying hemoglobin production per cell, which was done using HPLC to quantify the amount of heme released by hemoglobin based on a standard curve. This analysis revealed generally elevated hemoglobin production across *EPOR* (iEPOR)-edited clones cultured with BB compared to the same clones cultured with EPO (median of 33.1 vs. 24.4pg hemoglobin per cell, respectively; **Supplementary Fig. S11C** ). In contrast, clones edited with *PGK* (iEPOR) showed higher hemoglobin production when cultured with EPO (median of 21.1 vs. 32.1pg hemoglobin per cell with BB vs. EPO, respectively). Importantly, these levels of hemoglobin production are within the range expected for normal RBCs in the blood stream (25.4-34.6pg/cell)^31,32^. We note that while transfused RBCs typically produce more HbA than HbF, the healthy phenotype of patients with hereditary persistence of fetal hemoglobin (HPFH) and the recent approval of Casgevy to induce high levels of HbF to treat sickle cell disease and β-thalassemia provide support that a blood product with high HbF should be both safe and effective^33,34^.

## Discussion

In this work we combined synthetic protein engineering with the specificity of homology-directed repair genome editing to enable small molecule control of cell differentiation and behavior. By first optimizing highly effective small molecule-responsive receptors and then integrating them into endogenous regulatory machinery, we effectively recapitulated native receptor signaling. These efforts enable cell signaling to be stimulated by low-cost small molecules instead of recombinant cytokines currently required for *ex vivo* cell manufacturing. In this specific instance, EPO is one of the most expensive components of erythroid-promoting media^7^. Here, we demonstrate that *EPOR* (iEPOR)-edited cells cultured with a small molecule are capable of achieving equivalent erythroid differentiation, transcriptomic changes, and hemoglobin production compared to cells cultured with EPO. For comparison, we determined the cost per mg of the largest commercially available units of recombinant human EPO and AP20187 (BB) small molecule. We found that the price per mg of BB was nearly 50-fold less than that of recombinant EPO (**Fig. 5F** ). In addition, 1/10^th^ the amount of BB compared to recombinant EPO was required to yield equivalent erythroid production from *EPOR* (iEPOR)-edited iPSCs. Taken together, the corresponding estimates for cost of EPO required to produce a single unit of RBCs at a culture density of 5e7/mL is $1,246.50, conversely the cost to produce an equivalent amount of RBCs using BB is $2.25. While the tools and editing strategies defined in the work enable replacement of recombinant EPO with low-cost small molecule, additional key advances must be achieved for *ex vivo* RBC production to become biologically and economically feasible. These include further reducing the cost of erythroid-promoting media, better replicating high-density RBC production that occurs *in vivo*^30,35^, and improving enucleation of adult hemoglobin-producing RBCs^36^. This is a multi-faceted problem and will require sustained efforts to further reduce production expenses. Nevertheless, by eliminating the requirement of EPO cytokine in erythroid-promoting media, this work brings us one major step closer to establishing *ex vivo* RBC production as a scalable and renewable source of blood cells for transfusion medicine.

More broadly, we envision a future where clinically relevant cell types may be manufactured off-the-shelf and at scale to meet the broad spectrum of patient needs. However, significant advances are needed to improve affordability and accessibility to patients. Given the complexities of large-scale cell manufacturing, many innovations have been accomplished by mechanical engineers who have developed improved bioreactors^30,37,38^. Our work demonstrates how challenges within this space may also be addressed by genome engineers to create more effective producer cells to seed these advanced bioreactors. Because dependence on expensive cytokines is a common barrier to scalable production of any cells *ex vivo*, the strategies defined in this work may be readily adapted to enable large-scale production of platelets, neutrophils, T cells, and many other clinically relevant cell types. This will ensure that advancements in cell engineering may be rapidly translated to patients at a cost that is both affordable and accessible.

Finally, this work demonstrates the power of iterative design-build-test cycles to rapidly improve function of synthetic proteins. In this work, test cycle 1 defined the ideal placement of an FKBP domain within the EPO receptor. Test cycle 2 enhanced efficacy of iEPOR designs by incorporation of signal peptides and a naturally occurring *EPOR* mutation.

Finally, test cycle 3 defined the ideal expression profile of our optimized iEPOR cassette when placed under a variety of exogenous and endogenous promoters. Perhaps unsurprisingly, we find that integration of the optimized iEPOR at the endogenous *EPOR* locus best recapitulates native EPOR signaling, an engineering attribute enabled by homology-directed repair genome editing. In addition, given the incredible modularity of membrane-bound receptors^39^, it is possible that the small molecule-inducible architecture defined in this work may inform the design of other potentially useful small molecule-inducible receptors to modulate a wide variety of cell signals. If so, it is likely that synthetic receptor function may be fine-tuned by design-build-test cycles as well as genome editing to mediate precise integration into the genome. Gaining precisely regulated and tunable control over cells will thus pave the way for increasingly sophisticated cell engineering applications.

## Methods

### Integration vector design

Integration vectors were designed such that the left and right homology arms (LHA and RHA, respectively) are immediately flanking the cut site in exon 2 of the *CCR5* locus or exon 8 of the *EPOR* locus. For *HBA1* integration, full gene replacement was achieved as previously described^10^ using split homology arms—the LHA corresponding to the region immediately upstream of the start codon and RHA corresponding to the region immediately downstream of the cut site in the 3’ UTR of the *HBA1* gene. Homology arm length ranged from 400-1000bp. For FKBP-EPOR chimeras, flexible GGGGS linkers were added between FKBP domains and SPs and the *EPOR* gene. When placing the FKBP domain immediately adjacent to the EPOR transmembrane domain, the TM domain was defined as amino acid sequence PLILTLSLILVVILVLLTVLALLSH. EPOR SP was defined as amino acid sequence MDHLGASLWPQVGSLCLLLAGAAW. IL6 SP was defined as amino acid sequence MNSFSTSAFGPVAFSLGLLLVLPAAFPAP. The FKBP sequence used corresponded to amino acid sequence MLEGVQVETISPGDGRTFPKRGQTCVVHYTGMLEDGKKVDSSRDRNKPFKFMLGKQEVIRGWEEGVAQMSVGQRAKLTISPDYAYGATGHPGIIPPHATLVFDVELLKLE. Finally, to avoid the possibility of unintended recombination of iEPOR with the endogenous locus for *EPOR* (iEPOR)-edited conditions, we disguised homology of iEPOR by creating silent mutations within the *EPOR* domains at every possible codon, with a preference for codons that occurred more frequently throughout the human genome^40^. All custom sequences for cloning were ordered from Integrated DNA Technologies (IDT; Coralville, Iowa, USA). Gibson Assembly MasterMix (New England Biolabs, Ipswich, MA, USA) was used for the creation of each vector as per the manufacturer’s instructions.

### AAV6 DNA repair template production & purification

All AAV6 vectors were cloned into the pAAV-MCS plasmid (Agilent Technologies, Hayward, CA, USA), which contains inverted terminal repeats (ITRs) derived from AAV2. To produce AAV6 vectors, we seeded HEK293T cells (Life Technologies, South San Francisco, CA, USA) in 2-5 15cm^2^ dishes at 13-15×10^6^ cells per plate; 24h later, each dish was transfected using 112μg polyethyleneimine, 6μg of ITR-containing plasmid, and 22μg of pDGM6 (gift from D. Russell, University of Washington), which contains the AAV6 cap genes, AAV2 rep genes, and Ad5 helper genes. After 48-72h of incubation, cells were collected and AAV6 capsids were isolated using the AAVPro Purification Kit (All Serotypes, Takara Bio, San Jose, USA), as per the manufacturer’s instructions. AAV6 vectors were titered using a Bio-Rad QX200 ddPCR machine and QuantaSoft software (v.1.7, Bio-Rad, Hercules, CA, USA) to measure the number of vector genomes as described previously^10^.

### HSPC culture

Human CD34^+^ HSPCs were cultured as previously described^10^. CD34^+^ HSPCs were sourced from fresh cord blood (generously provided by the Stanford Binns Family Program for Cord Blood Research) and Plerixafor- and/or G-CSF-mobilized peripheral blood (AllCells, Alameda, CA, USA or STEMCELL Technologies, Vancouver, Canada). CD34^+^ HSPCs were cultured at 1-5×10^5^ cells/mL in StemSpan SFEMII (STEMCELL Technologies) or Good Manufacturing Practice Stem Cell Growth Medium (SCGM, CellGenix, Freiburg, Germany) base medium supplemented with a human cytokine (PeproTech, Rocky Hill, NJ, USA) cocktail: stem cell factor (100ng/mL), thrombopoietin (100ng/mL), Fms-like tyrosine kinase 3 ligand (100 ng/ml), interleukin-6 (100ng/mL), streptomycin (20mg/mL)(ThermoFisher Scientific, Waltham, MA, USA), and penicillin (20U/mL)(ThermoFisher Scientific, Waltham, MA, USA), and 35nM of UM171 (cat.:A89505; APExBIO, Houston, TX, USA). The cell incubator conditions were 37°C, 5% CO_2_, and 5% O_2_.

### Genome editing of HSPCs

Chemically modified CRISPR guide RNAs (gRNAs) used to edit CD34^+^ HSPCs at *CCR5* , *HBA1*, and *EPOR* were purchased from Synthego (Redwood City, CA, USA). The gRNA modifications added were 2’-O-methyl-3’-phosphorothioate at the three terminal nucleotides of the 5’ and 3’ ends, as described previously^41^. The target sequences for gRNAs were as follows: *CCR5*: 5’-GCAGCATAGTGAGCCCAGAA-3’; *HBA1*: 5’-GGCAAGAAGCATGGCCACCGAGG-3’; and *EPOR* : 5’-AGCTCAGGGCACAGTGTCCA-3’. All Cas9 protein was purchased from Aldevron (Alt-R S.p. Cas9 Nuclease V3; Fargo, ND, USA). Cas9 ribonucleoprotein (RNP) complexes were created at a Cas9/gRNA molar ratio of 1:2.5 at 25°C for a minimum of 10mins before electroporation. CD34^+^ cells were resuspended in P3 buffer plus supplement (cat.: V4XP-3032; Lonza Bioscience, Walkersville, MD, USA) with complexed RNPs and electroporated using the Lonza 4D Nucleofector (program DZ-100). Cells were plated at 1-2.5×10^5^ cells/mL following electroporation in the cytokine-supplemented media described previously. Immediately following electroporation, AAV6 was supplied to the cells at between 2.5-5e3 vector genomes per cell. The small molecule AZD-7648, a DNA-dependent protein kinase catalytic subunit inhibitor, was also added to cells immediately post-editing for 24h at 0.5nM to improve homology-directed repair frequencies as previously reported^42^.

### Ex vivo erythroid differentiation

Following editing, HSPCs derived from healthy patients or iPSC-derived HPCs were cultured for 14d at 37°C and 5% CO2 in SFEMII medium (STEMCELL Technologies, Vancouver, Canada) as previously described^19,20^. SFEMII base medium was supplemented with 100U/mL penicillin/streptomycin (ThermoFisher Scientific, Waltham, MA, USA), 10ng/mL stem cell factor (PeproTech, Rocky Hill, NJ, USA), 1ng/mL interleukin-3 (PeproTech, Rocky Hill, NJ, USA), 3U/mL EPO (eBiosciences, San Diego, CA, USA), 200μg/mL transferrin (Sigma-Aldrich, St. Louis, MO, USA), 3% antibody serum (heat-inactivated; Sigma-Aldrich), 2% human plasma (isolated from umbilical cord blood provided by Stanford Binns Cord Blood Program), 10μg/mL insulin (Sigma-Aldrich), and 3U/mL heparin (Sigma-Aldrich). In the first phase, at days 0–7 of differentiation (day 0 being 2-3d post editing), cells were cultured at 1×10^5^ cells/mL. In the second phase (days 7–10), cells were maintained at 1×10^5^ cells/mL and IL-3 was removed from the culture. In the third phase (days 11–14), cells were cultured at 1×10^6^ cells/mL and transferrin was increased to 1mg/mL. For -EPO conditions, cells were cultured in the same culture medium as listed above except for the removal of EPO from the media. For conditions with the addition of BB homodimerizer (AP20187)(Takara Bio, San Jose, USA), 1μL of 0.5mM BB was diluted in 999μL PBS (HI30; BD Biosciences, San Jose, CA, USA), of which 2μL of the dilution was added for every 1mL of differentiation media to reach a desired concentration of 1nM. Fresh BB was added at each media change (Day 0, 4, 7, 11). For experiments requiring additional dilutions, BB was diluted further in PBS to reach the required concentration (as low as 1pM).

### Immunophenotyping of differentiated erythrocytes

HSPCs subjected to erythroid differentiation were analyzed at d14 for erythrocyte lineage-specific markers using a FACS Aria II and FACS Diva software (v.8.0.3; BD Biosciences, San Jose, CA, USA). Edited and unedited cells were analyzed by flow cytometry using the following antibodies: hCD45 V450 (1:50 dilution; 2µl in 100µl of pelleted RBCs in 1×PBS buffer; HI30; BD Biosciences), CD34 APC (1:50 dilution; 561; BioLegend, San Diego, CA, USA), CD71 PE-Cy7 (1:500 dilution; OKT9; Affymetrix, Santa Clara, CA, USA), and CD235a PE (GPA)(1:500 dilution; GA-R2; BD Biosciences). In addition to cell-specific markers, cells were also stained with Ghost Dye Red 780 (Tonbo Biosciences, San Diego, CA, USA) to measure viability.

### Editing frequency analysis

Between 2-4d post editing, HSPCs were harvested and QuickExtract DNA extraction solution (Epicentre, Madison, WI, USA) was used to collect genomic DNA (gDNA). Additional samples were collected at various stages of differentiation (d4, 7, 11, and 14 of erythroid differentiation) gDNA was then digested using BamH1-HF as per the manufacturer’s instructions (New England Biolabs, Ipswich, MA, USA). Percentage of targeted alleles within a cell population was measured with a Bio-Rad QX200 ddPCR machine and QuantaSoft software (v.1.7; Bio-Rad, Hercules, CA, USA) using the following reaction mixture: 1-4μL of digested gDNA input, 10μL of ddPCR SuperMix for Probes (no dUTP)(Bio-Rad), primer/probes (1:3.6 ratio; Integrated DNA Technologies, Coralville, IA, USA) and volume up to 20μL with H2O. ddPCR droplets were then generated following the manufacturer’s instructions (Bio-Rad): 20μL of ddPCR reaction, 70μL of droplet generation oil, and 40μL of droplet sample. Thermocycler (Bio-Rad) settings were as follows: 98°C (10mins), 94°C (30s), 57.3°C (30s), 72°C (1.75mins)(return to step 2×40-50 cycles), and 98°C (10mins). Analysis of droplet samples was performed using the QX200 Droplet Digital PCR System (Bio-Rad). To determine percentages of alleles targeted, the numbers of Poisson-corrected integrant copies/mL were divided by the numbers of reference DNA copies/mL. The following primers and 6-FAM/ZEN/IBFQ-labeled hydrolysis probes were purchased as custom-designed PrimeTime quantitative PCR (qPCR) assays from Integrated DNA Technologies: All *HBA1* vectors: forward: 5’-AGTCCAAGCTGAGCAAAGA-3’, reverse: 5’-ATCACAAACGCAGGCAGAG-3’, probe: 5’-CGAGAAGCGCGATCACATGGTCCTGC-3’; all *CCR5* vectors: forward: 5’-GGGAGGATTGGGAAGACAAT-3’, reverse: 5’-TGTAGGGAGCCCAGAAGAGA-3’, probe: 5’-CACAGGGCTGTGAGGCTTAT-3’. The primers and HEX/ZEN/IBFQ-labeled hydrolysis probe, purchased as custom-designed PrimeTime qPCR Assays from Integrated DNA Technologies, were used to amplify the *CCRL2* reference gene: forward: 5’-GCTGTATGAATCCAGGTCC-3’, reverse: 5’-CCTCCTGGCTGAGAAAAAG-3’, probe: 5’-TGTTTCCTCCAGGATAAGGCAGCTGT-3’.

### AlphaFold2 structural predictions

Energy-predicted structures were derived by applying AlphaFold2 (v2.3.2)^21^ on wild-type EPOR, truncated EPOR, and iEPOR sequences. Five differently trained neural networks were applied to produce unrelaxed structure predictions. Energy minimization was applied to the best predicted unrelaxed structure (highest average highest average predicted distance difference test (pLDDT) and lowest predicted aligned error) to produce the optimal relaxed structure.

### Hemoglobin tetramer analysis

Frozen pellets of approximately 1×10^6^ cells *ex vivo* -differentiated erythroid cells were thawed and lysed in 30μL of RIPA buffer with 1x Halt Protease Inhibitor Cocktail (ThermoFisher Scientific, Waltham, MA, USA) for 5mins on ice. The mixture was vigorously vortexed and cell debris was removed by centrifugation at 13,000 RPM for 10mins at 4°C. HPLC analysis of hemoglobins in their native form was performed on a cation-exchange PolyCAT A column (35 × 4.6mm^2^, 3µm, 1,500Å)(PolyLC Inc., Columbia, MD, USA) using a Perkin-Elmer Flexar HPLC system (Perkin-Elmer, Waltham, MA, USA) at room temperature and detection at 415nm. Mobile phase A consisted of 20mM Bis-tris and 2mM KCN at pH 6.94, adjusted with HCl. Mobile phase B consisted of 20mM Bis-tris, 2mM KCN, and 200mM NaCl at pH 6.55. Hemolysate was diluted in buffer A prior to injection of 20μL onto the column with 8% buffer B and eluted at a flow rate of 2 mL/min with a gradient made to 40% B in 6mins, increased to 100% B in 1.5mins, returned to 8% B in 1min, and equilibrated for 3.5mins. Quantification of the area under the curve of peaks was performed with TotalChrom software (Perkin-Elmer) and raw values were exported to GraphPad Prism software (v9) for plotting and further analysis.

### Bulk RNA-sequencing

Total RNA was extracted from frozen pellets of approximately 1×10^6^ cells per condition using RNeasy Plus Micro Kit (Qiagen, Redwood City, CA, USA) according to the manufacturer’s instructions. Sequencing was provided by Novogene (Sacramento, CA, USA) and raw FASTQ files were aligned to the GRCh38 reference genome extended with the iEPOR target sequence and quantified using Salmon (v1.9.0)^43^ with default parameters. Quality control was performed by Novogene.

### Differential gene expression analysis & gene set enrichment analysis

The estimated gene expression counts were used with DESeq2^44^ to conduct differential gene expression analysis between sample groups. Mitochondrial and lowly expressed genes were removed (sum NumReads <1). The top 50 up- and down-regulated genes based on adjusted p-value were isolated and analyzed with Enrichr^45^ to yield functional annotations.

### Principal component analysis & gene distribution plots

Mitochondrial genes were removed from the gene expression matrix (TPM) and the remaining genes were used to conduct principal component analysis with all samples. Gene expression for experimental and control groups were averaged and log-normalized. Average gene expression distributions were plotted using Seaborn (https://github.com/atsumiando/RNAseq_figure_plotter_python).

### Gene co-expression network analysis

The TPM-normalized gene expression matrix of all *PGK* (iEPOR)-, *HBA1*(iEPOR)-, and *EPOR* (iEPOR)-edited conditions (n=10) was used to construct a pairwise gene similarity matrix where each entry represented the Spearman correlation coefficient between a pair of genes. The correlation between a specified set of *EPOR* -related genes was compared for both *EPOR* and *iEPOR* to determine which genes *iEPOR* adequately mimics in the immediate gene co-expression network of native *EPOR* .

### iPSC line & culture

A previously published iPSC line, PB005 derived from peripheral blood of a donor with O^-^ blood type was used in this study^46^. iPSCs were cultured and maintained in mTeSR1 medium (cat.: 85850; STEMCELL Technologies, Vancouver, Canada) on Matrigel (cat.: 354277; Corning, NY, USA)-coated plates. For passaging, cells at a confluency of 80-90% were incubated with Accutase (cat.: AT104; Innovative Cell Technologies, San Diego, USA) for 5-7mins to dissociate into single cells and replated in mTeSR1 medium supplemented with 10mM of ROCKi (Y27632; cat.: 10005583; Cayman Chemical, Ann Arbor, MI, USA). After 24h, cells were maintained in fresh mTeSR1 medium with daily media changes. For freezing iPS cells, STEM-CELLBANKER freezing medium (cat.: 11924; Amsbio, Cambridge, MA, USA) was used.

### Genome editing of iPSCs

iPSCs were genome edited using the CRISPR/AAV platform as described previously^17,42^. Cas9 RNP complex was formed by combining 5μg of Cas9 (Alt-R S.p. Cas9 Nuclease V3; Fargo, ND, USA) and 2μg of gRNA (Synthego, Redwood City, CA, USA) and incubating at room temperature for 15mins. iPSCs pre-treated with ROCKi (Y27632; cat.: 10005583; Cayman Chemical, Ann Arbor, MI, USA) for 24 hours were dissociated with Accutase (cat.: AT104; Innovative Cell Technologies, San Diego, USA) into single cells. 1-5x10^5^ iPSCs were resuspended in 20μL of P3 primary cell nucleofector solution plus supplement (cat.: V4XP-3032; Lonza Bioscience, Walkersville, MD, USA) along with the RNP complex and nucleofected using Lonza 4D Nucleofector (program CA-137). After nucleofection, iPSCs were plated in mTeSR1 medium supplemented with ROCKi, 0.25μM AZD7648 (cat.: S8843; Selleck Chemicals, Houston, TX, USA) and AAV6 donor at 2.5x10^3^ vector genomes per cell, based on ddPCR titers as above. After 24h, cells were switched to medium with mTeSR1 and ROCKi. From the following day, cells were maintained in mTeSR1 medium without ROCKi.

### Single-cell cloning of iPSCs

To isolate single cell clones, genome-edited iPSCs were plated at a density of 250 cells per well of a 6-well plate in mTeSR1 medium supplemented with 1x CloneR2 reagent (cat.: 100-0691; STEMCELL Technologies, Vancouver, Canada). After 48h, cells were switched to fresh mTeSR1 medium with 1x CloneR2 and incubated for 2d. Following this, iPSCs were maintained in mTeSR1 medium without CloneR2 with daily media changes. At d7-10, single cell colonies were picked by scraping and propagated individually. The isolated single cell iPSCs were genotyped using PCR with primers annealing outside the homology arms to identify clones with bi-allelic knock-in. The following primers were used for genotyping: *CCR5* integration: forward: 5’-CTCATAGTGCATGTTCTTTGTGGGC-3’, reverse: 5’-CCAGCCCAGGCTGTGTATGAAA-3’; *EPOR* integration: forward: 5’-GCCACATGGCTAGAGTGGTAT-3’, reverse: 5’-CTTTCTTAGAACATGGCCTGATTCAGA-3’.

### iPSC-to-erythrocyte differentiation

iPSCs were differentiated into CD34^+^ HPCs using the STEMdiff Hematopoietic Kit (cat.: 05310; STEMCELL Technologies, Vancouver, Canada) according to the manufacturer’s protocol. Briefly, iPSCs at 70-80% confluency were dissociated into aggregates using ReLeSR (cat.: 100-0484; STEMCELL Technologies). Aggregates were then diluted 10-fold, and 100μL of the diluted suspension was aliquoted into a 96-well plate for quantification. Approximately 80 aggregate colonies were subsequently plated per well of a 12-well plate pre-coated with Matrigel and maintained in mTeSR1 medium. 24h post-plating, the number of colonies per well was manually quantified, and the medium was replaced with differentiation medium A. The medium was then changed according to the kit’s instructions for a total of 12 days. On d12, suspension cells were harvested by pipetting cells up and down to ensure a homogeneous cell suspension. To assess the efficiency of differentiation, as determined by CD34^+^/CD45^+^ expression, cells were analyzed using flow cytometry with the erythrocyte flow panel previously described for HSPCs. Following this, CD34^+^ cells were further differentiated into erythroid cells using the three-phase system described above, either in the presence or absence of EPO and BB.

### Heme detection analysis

Quantification of the amount of hemoglobin produced in cells was obtained by quantitative detection of the heme peak released from hemoglobin. Lysate were obtained from 1-2x10^5^ cells as frozen pellets, as described for hemoglobin tetramer analysis. The relationship between heme and hemoglobin was established from serially diluted hemolysate made with a blood sample of a known hemoglobin content. Detection of heme was performed by reverse-phase PerkinElmer Flexar HPLC system (PerkinElmer) with a Symmetry C18 column (4.6 ×^75mm, 3.5µm; Waters Corporation, Milford, MA, USA) at 415nm. Mobile phase^ A consisted of 10% methanol made in acetonitrile and mobile B of 0.5% trifluoroacetic acid in water adjusted at pH 2.9 with NaOH. Samples were injected at a flow rate of 2mL/min in 49% A, followed by a 3min gradient to 100% A. The column was then equilibrated to 49% A for 3mins.

### Statistical analysis

All statistical tests on experimental groups were done using GraphPad Prism software (v9). The exact statistical tests used for each comparison are noted in the individual figure legends.

## Data availability

RNA-seq data will be uploaded to the NCBI Sequence Read Archive submission. The filtered data for all figures in this study are provided in the Extended Data.

## Acknowledgements

The authors thank the following funding sources that made this work possible: Stanford Medical Scholars Research Program, the American Society of Hematology Minority Medical Student Award Program, and the Stanford Medical Scientist Training Program to S.E.L.; National Science Foundation Graduate Research Fellowship Program to B.J.L.; University of California, San Francisco NIH T32 Research Training in Transplant Surgery Fellowship to S.N.C.; University of California, San Francisco Program for Breakthrough Biomedical Research: New Frontier Research Award to M.K.C; and Mary Anne Koda-Kimble Seed Award for Innovation to M.K.C. We also would also like to thank the Stanford Binns Program for Cord Blood Research for providing CD34^+^ HSPCs and the FACS Core Facility at the Stanford Institute of Stem Cell Biology and Regenerative Medicine as well as University of California, San Francisco Flow Core for access to flow cytometry machines. Finally, we would like to thank Caleb Grossman for helpful discussions and feedback as we planned initial experiments.

