## Supplemental Figures for "Engineering inducible signaling receptors to enable erythropoietin-free erythropoiesis"

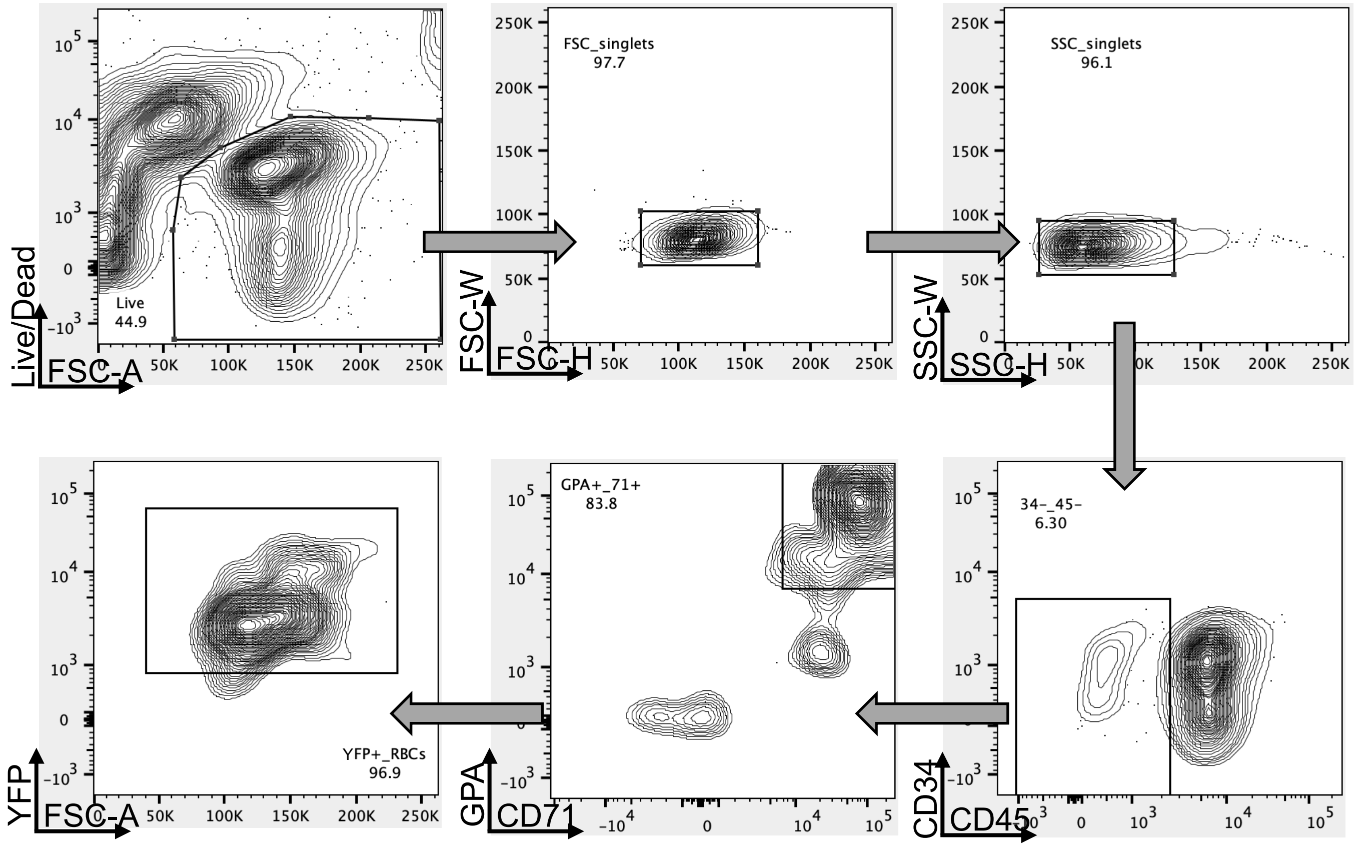


**Figure S1: Representative flow cytometry staining and gating scheme for erythroid differentiation analysis.** HSPCs were edited with iEPOR 1.5, subjected to *ex vivo* HSPC-to-erythroid cell differentiation in absence of EPO but presence of BB, and analyzed at day 14. Arrows indicate that only gated cells are displayed on the subsequent plot.

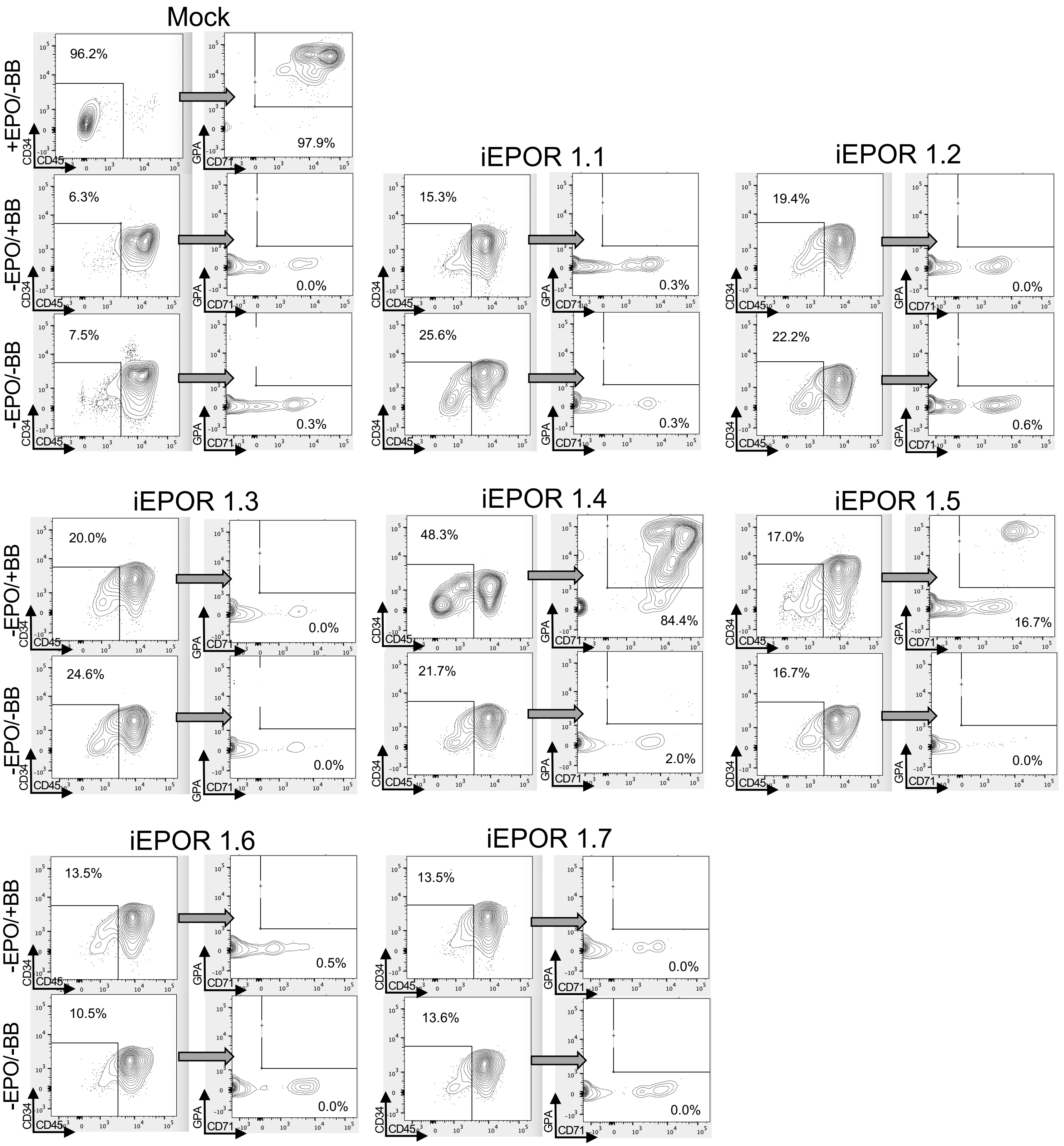


**Figure S2: Flow cytometry analysis of first-generation iEPORs.**
Representative flow cytometry staining and gating scheme for first-generation iEPOR-edited HSPCs at d14 of erythroid differentiation. Arrows indicate that only gated cells are displayed on the subsequent plot.

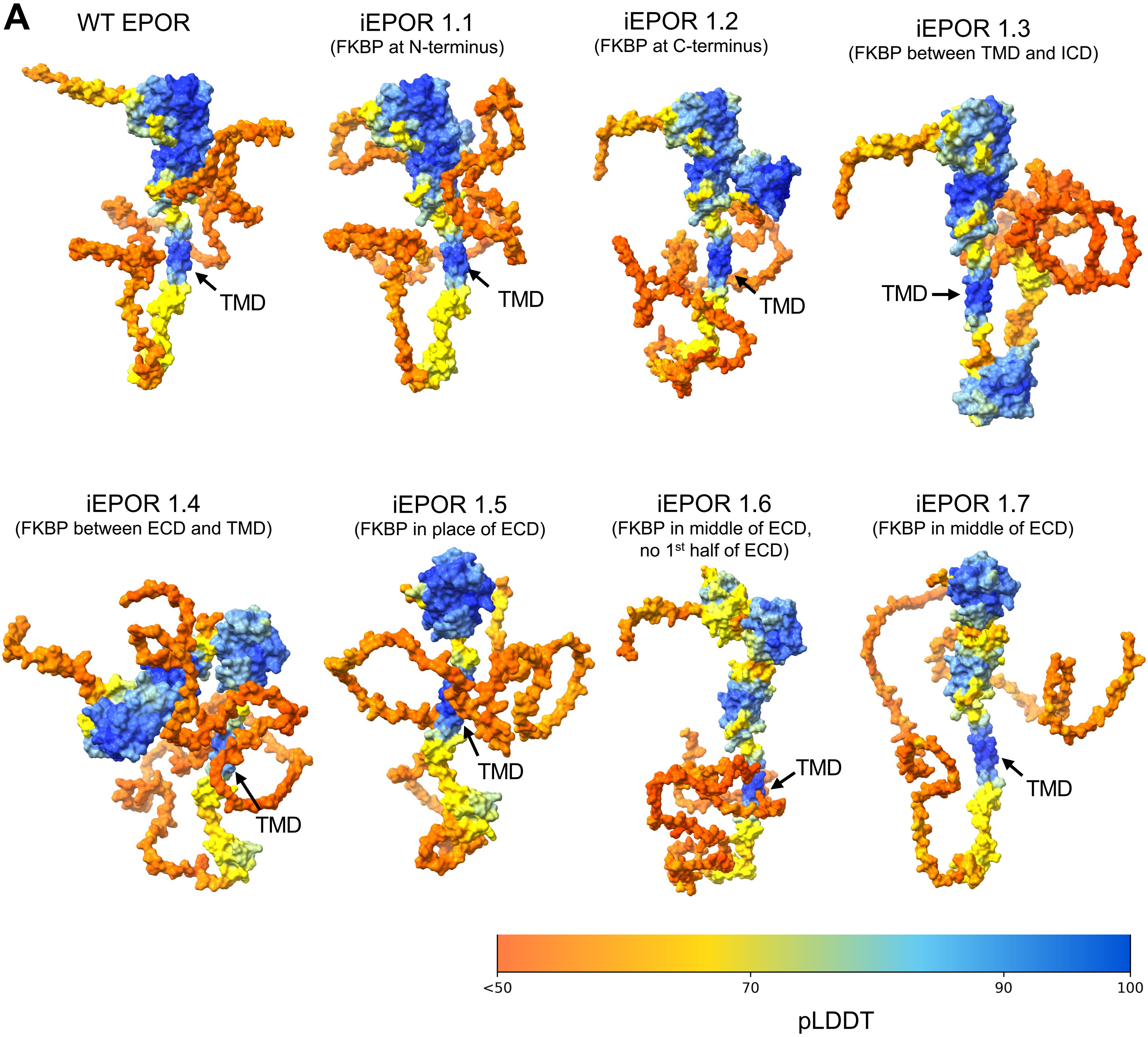


**Figure S3: *In silico* structure prediction of first-generation iEPORs.**
A: AlphaFold2-based structure prediction of wild-type (WT) EPOR and all candidate iEPORs. As a measure of structure confidence, predicted distance difference test (pLDDT) is displayed according to the color bar. Abbreviations as follows: ECD = extracellular domain; TMD = transmembrane domain; and ICD = intracellular domain. TMD labeled with an arrow as a reference point.

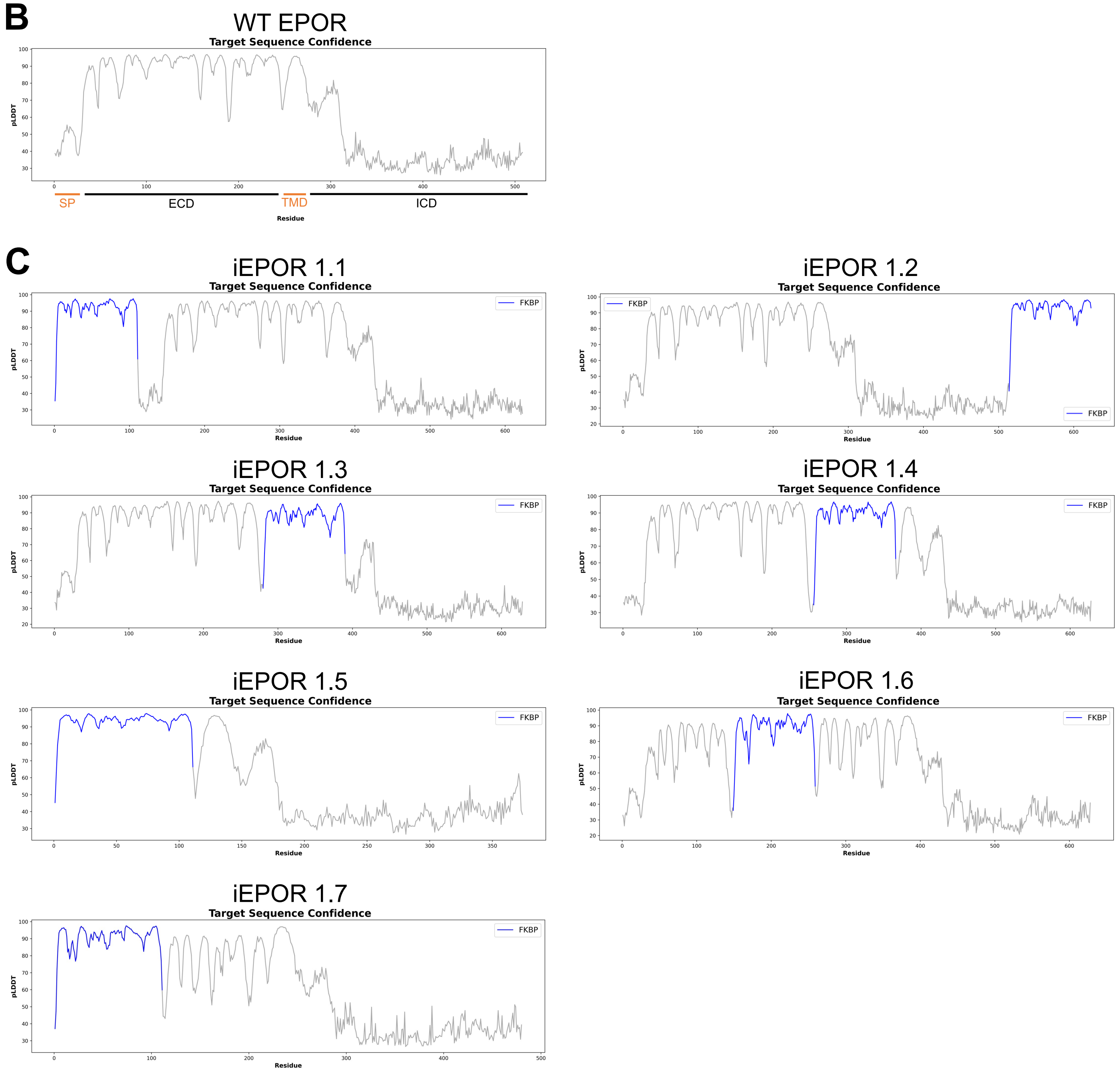


**Figure S3 (cont'd): *In silico* structure prediction of first-generation iEPORs.**
B: Plot of pLDDT of AlphaFold2-predicted structure for WT EPOR shown in Figure S3A. Annotations are as follows: SP = signal peptide; ECD = extracellular domain; TMD = transmembrane domain; and ICD = intracellular domain.
C: Plots of pLDDT of predicted structures for all candidate iEPORs shown in Figure S3A. FKBP domain is highlighted in blue.

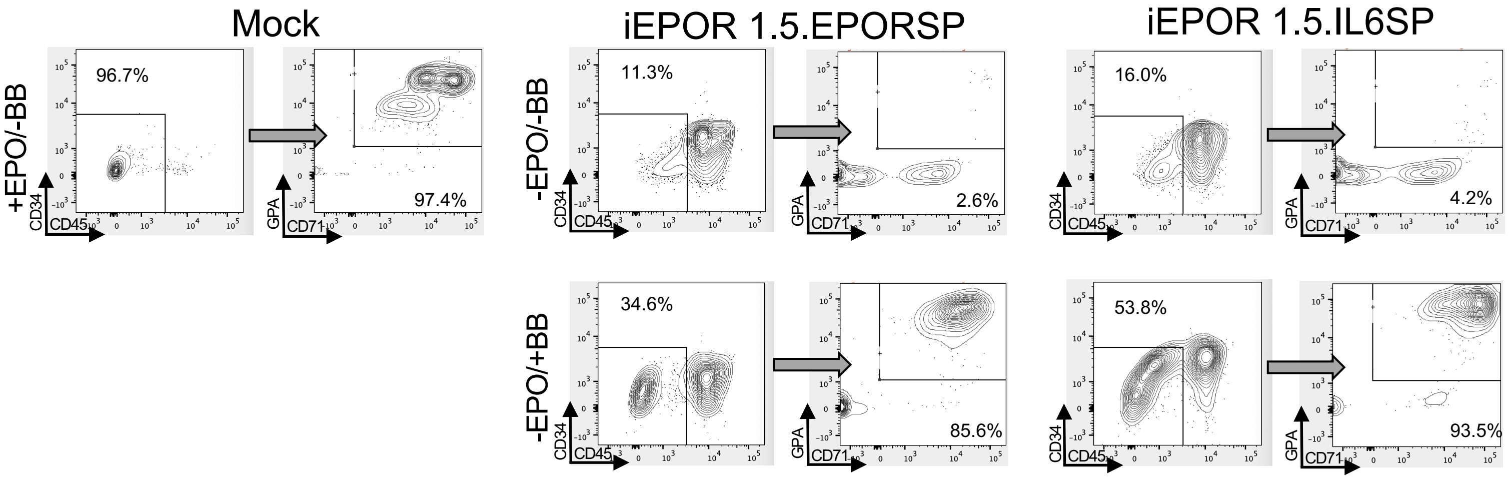


**Figure S4: Flow cytometry analysis of second-generation iEPORs.**
Representative flow cytometry staining and gating scheme for second-generation iEPOR-edited HSPCs at d14 of erythroid differentiation. Arrows indicate that only gated cells are displayed on the subsequent plot.

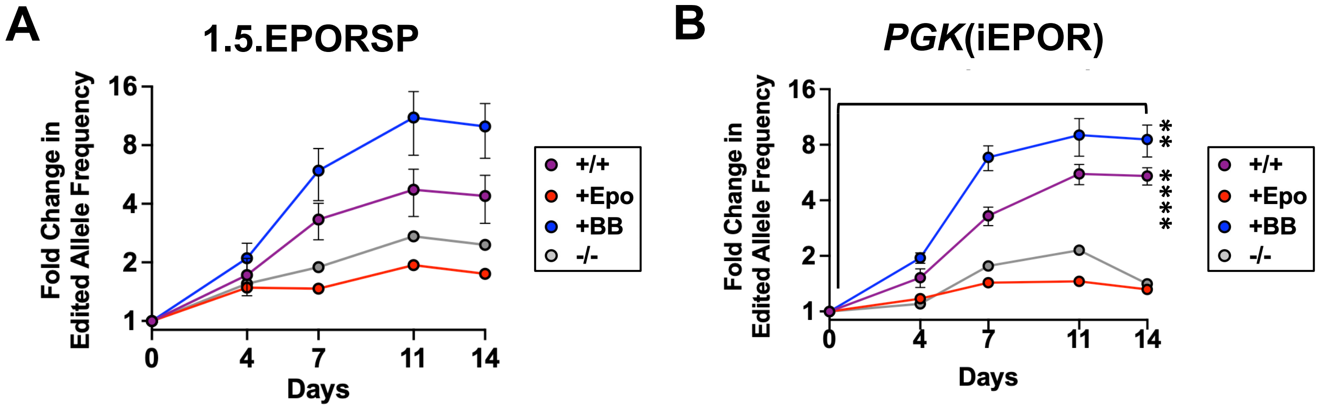


**Figure S5: Enrichment of edited cells over course of erythroid differentiation.**

A: Percent alleles edited with 1.5EPORSP measured by ddPCR over the course of differentiation +/-EPO and +/-BB. Bars represent median +/-SEM.

B: Percent alleles edited with *PGK*(iEPOR) measured by ddPCR over the course of differentiation +/-EPO and +/-BB. Bars represent median +/-SEM; ** = p<0.01 and **** = p<0.0001 comparing treatment at d0 vs. d14 by unpaired t-test.

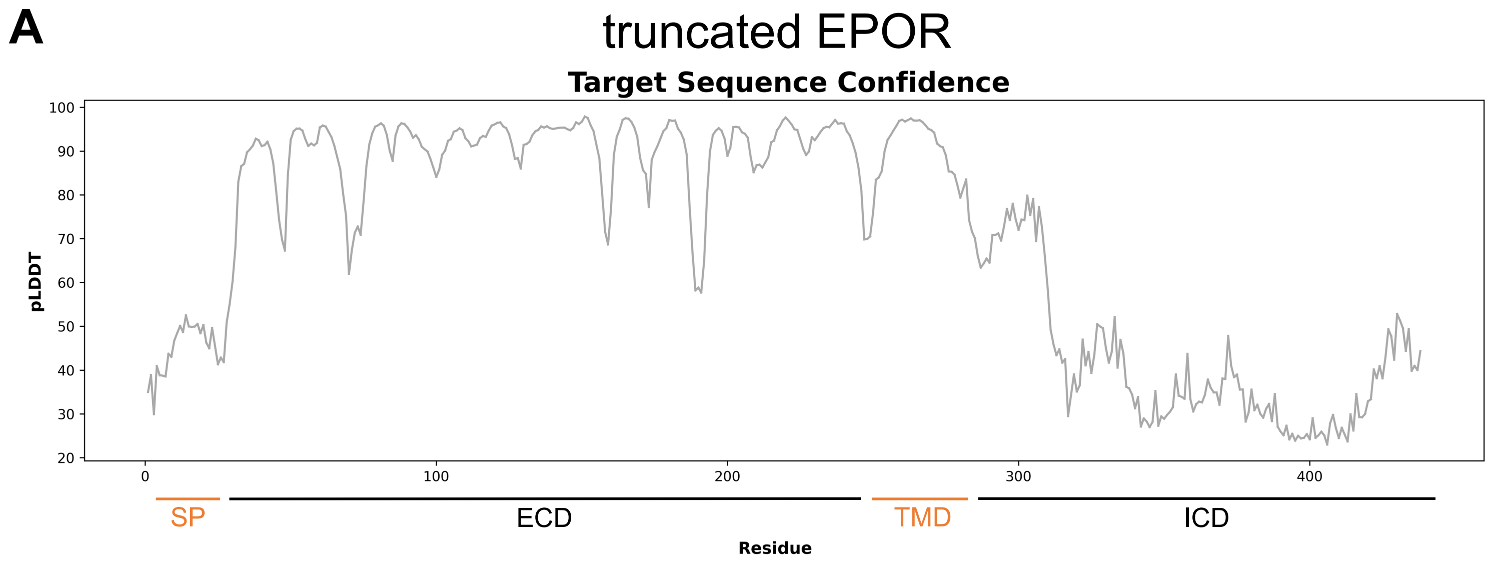


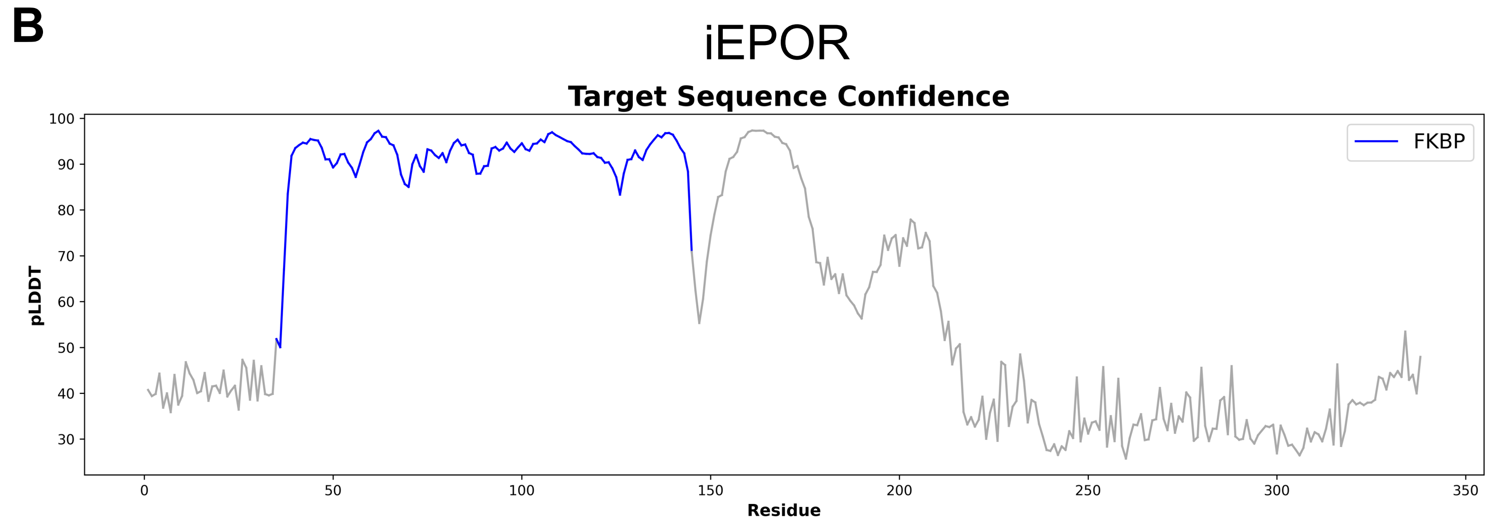


**Figure S6: *In silico* structure prediction of final iEPOR design.**
A: Plot of pLDDT of AlphaFold2-predicted structure for truncated EPOR shown in Fig. 2F. Annotations are as follows: SP = signal peptide; ECD = extracellular domain; TMD = transmembrane domain; and ICD = intracellular domain.
B: Plot of pLDDT of predicted structure for the optimized iEPOR shown in Fig. 2F. FKBP domain is highlighted in blue.

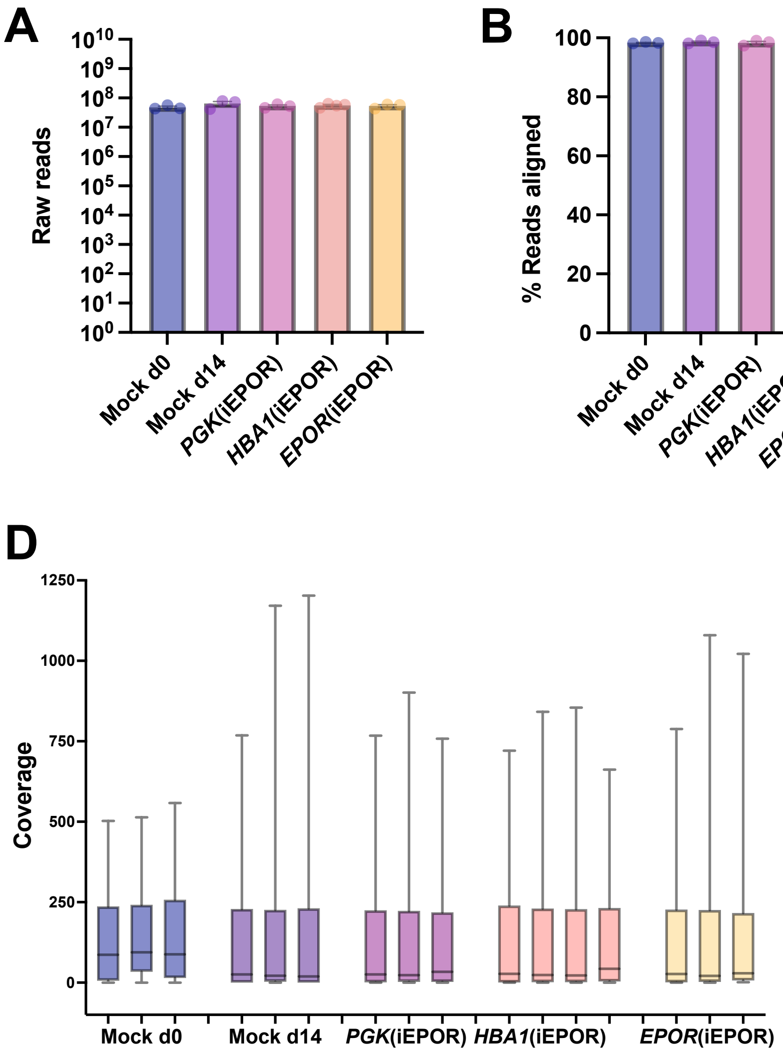


**Figure S7: RNA-sequencing quality.**

A: Summary of reads by condition.

B: Percentage of reads aligned to genome by condition.

C: Percentage of reads with Quality Score $\geq$20 by condition.

D: Coverage after normalization by condition. Box represents 25$^{th}$, 50$^{th}$, and 75$^{th}$ percentiles. Whiskers represent 10$^{th}$ and 90$^{th}$ percentiles.

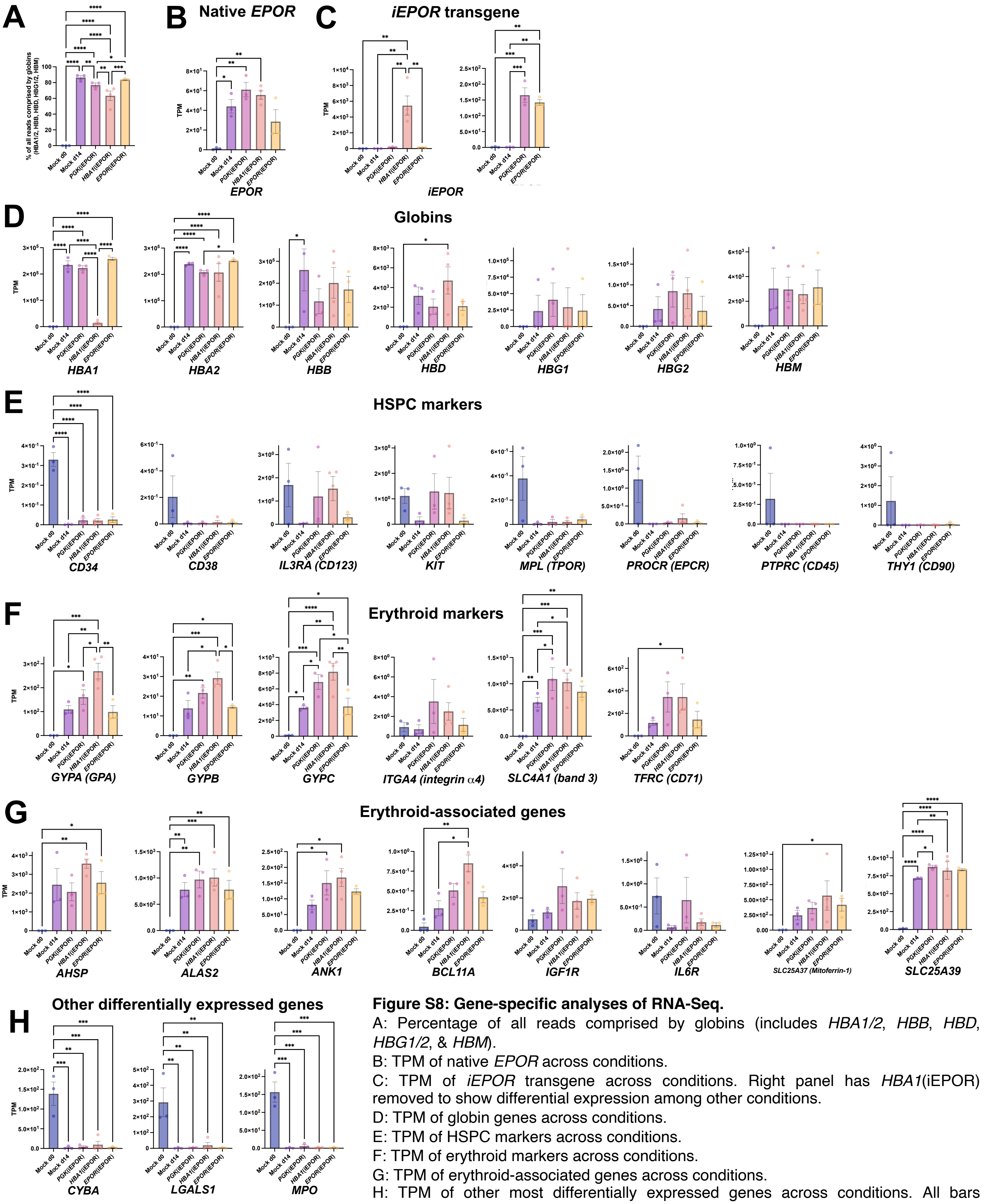


**Figure S8: Gene-specific analyses of RNA-Seq.**

A: Percentage of all reads comprised by globins (includes *HBA1/2*, *HBB*, *HBD*, *HBG1/2*, & *HBM*).

B: TPM of native *EPOR* across conditions.

C: TPM of *iEPOR* transgene across conditions. Right panel has *HBA1*(iEPOR) removed to show differential expression among other conditions.

D: TPM of globin genes across conditions.

E: TPM of HSPC markers across conditions.

F: TPM of erythroid markers across conditions.

G: TPM of erythroid-associated genes across conditions.

H: TPM of other most differentially expressed genes across conditions. All bars represent median +/-SEM; * = p<0.05, ** = p<0.01, *** = p<0.001, and **** = p<0.0001 by 2-way ANOVA for multiple comparisons.

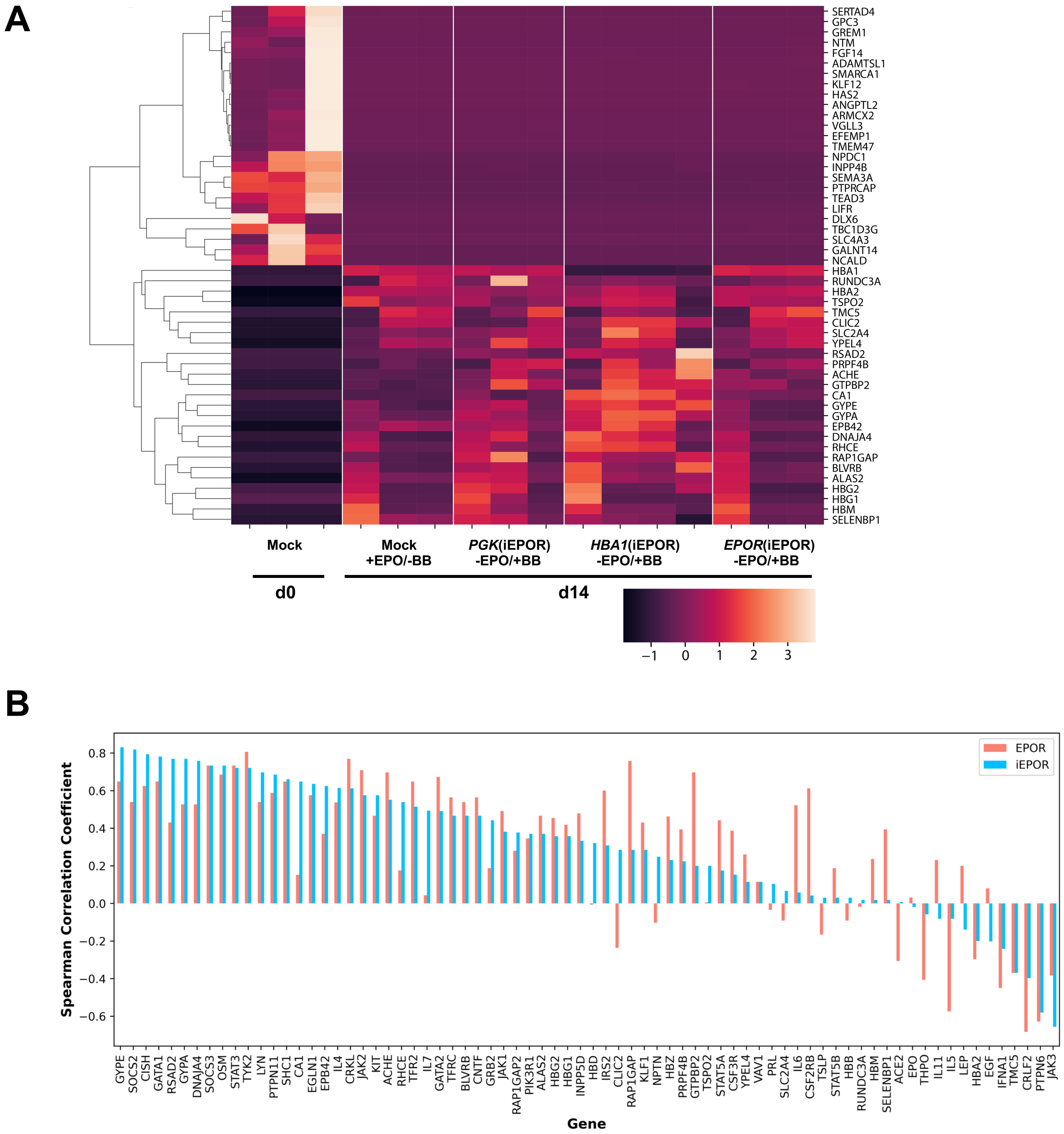


**Figure S9: Additional analyses of transcriptomic data.**

A: Heatmap plotting top 25 differentially upregulated and downregulated genes (comparing d0 vs. all d14 samples).

B: Spearman correlation coefficient is plotted for a specified set of EPOR-related genes to depict which genes iEPOR adequately mimics in the immediate gene co-expression network of native EPOR.

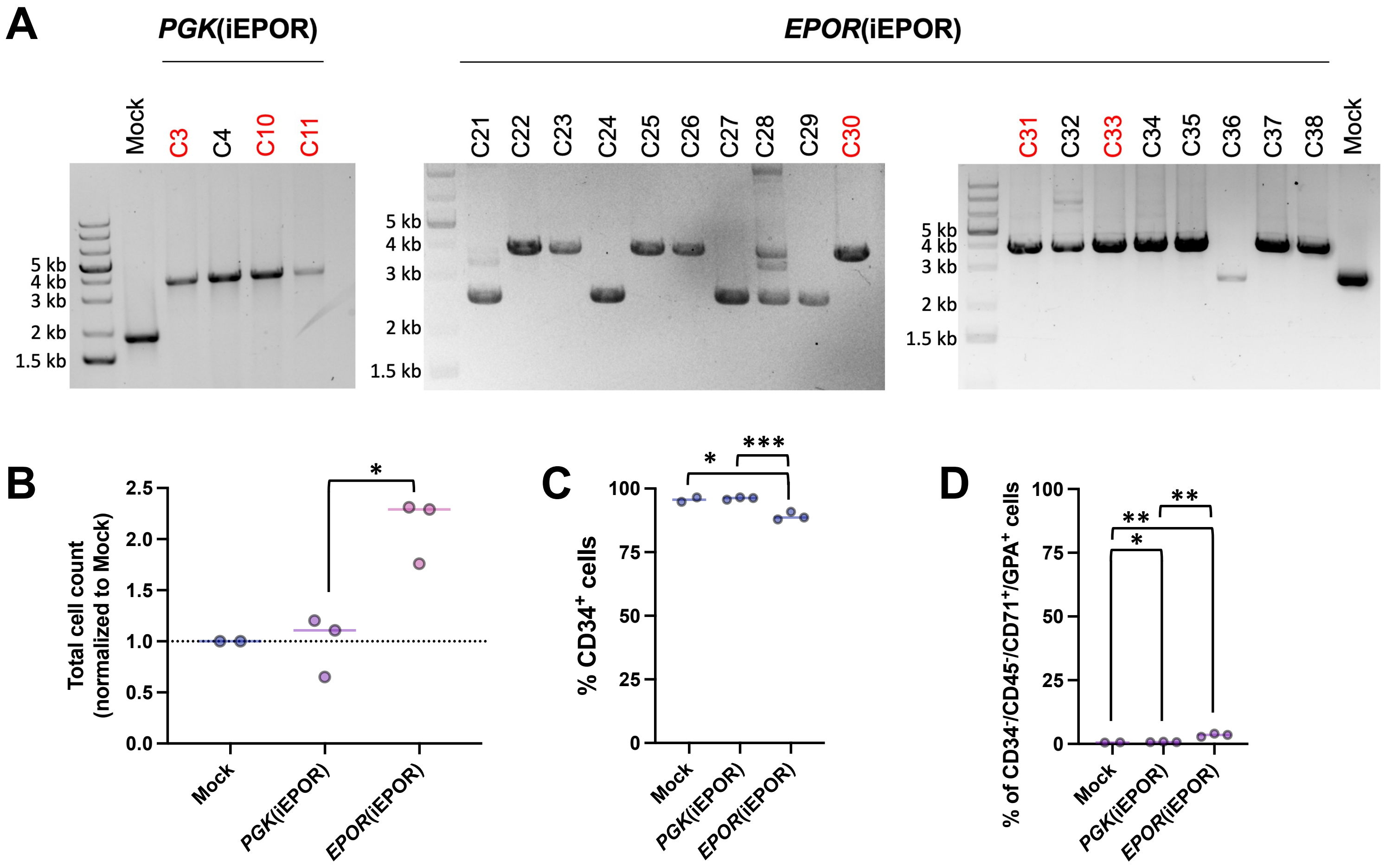


**Figure S10: Isolation of homozygous iEPOR-edited iPSC clones & downstream HPC differentiation.**

A: Agarose gel electrophoresis images of PCR-amplified genomic DNA surrounding the predicted *CCR5* and *EPOR* integration sites which was used to genotype iEPOR-edited iPSC clones. Homozygous clones chosen for downstream analysis are highlighted in red.

B: Total cell counts at the end of iPSC-to-HPC differentiation normalized to unedited cells. * = $p<0.05$ by unpaired t-test.

C: Percentage of cells staining $CD34^+$ by flow cytometry following iPSC-to-HPC differentiation. Bars represent median; * = $p<0.05$; *** = $p<0.001$ by unpaired t-test.

D: Percentage of cells that acquired erythroid markers ($CD34^-/CD45^-/CD71^+/GPA^+$) following iPSC-to-HPC differentiation. Bars represent median; * = $p<0.05$; ** = $p<0.01$ by unpaired t-test.

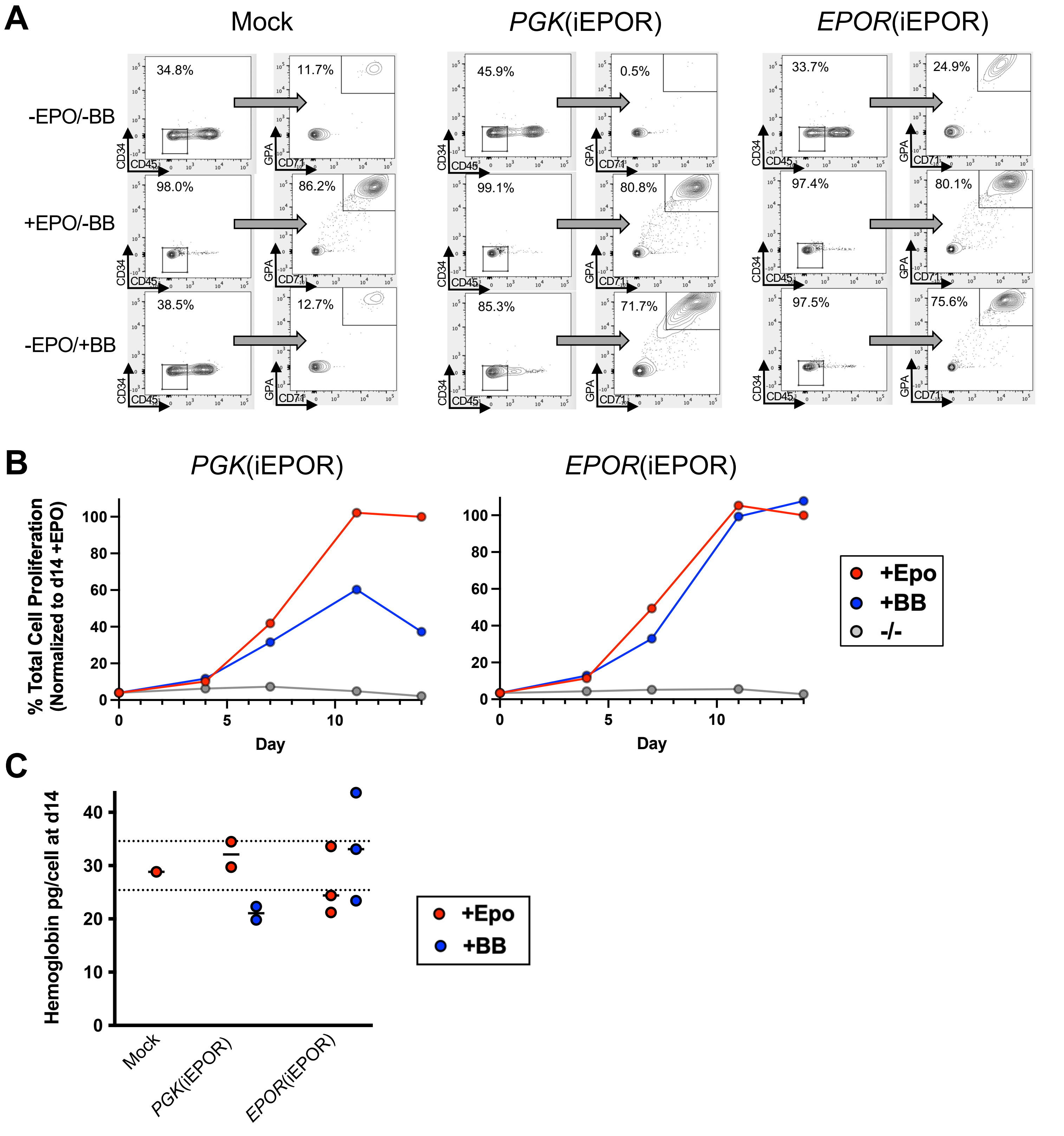


**Figure S11: Additional analyses of iEPOR-edited iPSC-derived erythroid cells.**

A: Representative flow cytometry staining and gating scheme for iEPOR-edited iPSCs at end of erythroid differentiation. Arrows indicate that only gated cells are displayed on the subsequent plot.

B: Percent of total cell proliferation normalized to clones cultured +EPO over the course of differentiation. Only the clone with greatest proliferation in presence of BB is shown in each editing condition. Bars represent mean +/SEM.

C: Hemoglobin production per cell for edited and unedited iPSCs at end of erythroid differentiation. Bars depict median value; dotted lines depict normal range of mean corpuscular hemoglobin concentration in peripheral blood.
